# New methods on the block: Taxonomic identification of archaeological bones in resin-embedded sediments through palaeoproteomics

**DOI:** 10.1101/2025.03.14.643068

**Authors:** Zandra Fagernäs, Gaudry Troché, Paul Goldberg, Jean-Jacques Hublin, Shannon P. McPherron, William Chase Murphree, Jesper V. Olsen, Dennis Sandgathe, Nikolay Sirakov, Marie Soressi, Tsenka Tsanova, Alain Turq, Michael Wierer, Frido Welker, Vera Aldeias

## Abstract

The integration of biomolecular studies of past organisms with geoarchaeological studies can significantly improve our understanding of the relative chronology and context of archaeologically (in)visible behaviours. However, the complexity and sedimentological heterogeneity of archaeological deposits at a microscopic scale is often not taken into consideration in biomolecular studies. Here, we investigate the preservation and retrieval of palaeoproteomic data from bone fragments embedded in Pleistocene resin-impregnated sediment blocks. We show that resin impregnation has minimal effect on skeletal protein taxonomic identifications in modern skeletal material, but observe an increase in oxidation-related post-translational modifications. We then successfully retrieve proteins from resin-impregnated blocks from the Palaeolithic sites of Bacho Kiro Cave, La Ferrassie and Quinçay. The taxonomic identifications of minute bones encased in resin are in line with previous analyses of the faunal communities of these sites, with a diversity of taxa (*Bos* sp./*Bison* sp., *Equus* sp., *Ursus* sp., and Caprinae) observed at a microscale in Bacho Kiro. This differs from results from La Ferrassie where most of the samples are identified as a single taxon (*Bos* sp./*Bison* sp.) across different areas of the site. The block from Quinçay only provided taxonomic identification of two out of eleven bone-derived samples, likely due to diagenesis. Our work indicates that palaeoproteomes can be retrieved from bone fragments at a microstratigraphic resolution, enabling the detailed study of faunal community composition at a scale that more closely matches that of past human occupations.

**Significance Statement:** Resin-embedded sediment blocks are widely used in archaeology and soil sciences to reconstruct past environments and human behavior, but their potential for biomolecular analysis is underexplored. Here, we demonstrate that ancient proteins can be successfully retrieved from bone fragments embedded in resin-impregnated sediment blocks from Pleistocene archaeological sites. Our findings show that resin impregnation has a minimal impact on protein recovery and that palaeoproteomics enables taxonomic identification of Pleistocene bone fragments at a microstratigraphic scale. This approach allows for reconstructing past faunal communities with unprecedented detail, improving our understanding of ancient ecosystems and the environmental contexts of early hominin occupations.

## Introduction

Archaeological sediments are excellent archives to study past environmental diversity. Therefore, integrating sedimentary approaches with biomolecular studies of past organisms can significantly improve understanding of the relative chronology and context of biomolecular finds (1, 2). Resin-impregnated sediment blocks are of interest for biomolecular archaeology, as they provide a high-resolution image of the past through the sampling of structurally intact (i.e., not excavated) components directly associated with sedimentary microstratigraphy. Lipid biomarkers have been found to be preserved in sediment blocks (3), opening up the possibility for fine-scale reconstruction of palaeoenvironments. Targeted microsampling of bone fragments and coprolites visually identified in resin-impregnated sediment blocks has previously been shown to lead to successful palaeogenetic analysis (4). Additionally, Massilani et al. found that resin impregnation had a negligible effect on DNA preservation and on success of DNA extraction, indicating that resin-impregnated blocks can be excellent sources for the study of the faunal community at archaeological sites in microstratigraphic context.

Palaeoproteomic analysis is commonly used to identify the taxonomy of fragmented bones, which do not preserve morphological features that can be used for species identification. Methods such as ZooMS (zooarchaeology by mass spectrometry (5)) and SPIN (species by proteome investigation (6)) have been developed to efficiently identify differences in preserved amino acid sequences between archaeological skeletal specimens and to assign them a taxonomic identity by comparison to databases of known protein sequences. These approaches can be especially useful when studying Pleistocene sites, as faunal assemblages of such antiquity can be highly fragmented and are challenging to identify by morphology (7). Additionally, they can lead to the identification of new hominin remains, which may otherwise not have been found due to the highly fragmented nature of the bones (8–11). However, the time depth of Pleistocene deposits implies that geological layers formed over thousands of years and are, likely, often palimpsests that encompass multiple hominin occupations and/or separate faunal accumulations not directly related to hominin activity. Relevant behavioural and environmental information encapsulated in such deposits can really only be captured at a microstratigraphic temporal resolution.

Here, we explore the potential of applying palaeoproteomics to taxonomically identify minute bone fragments within resin-impregnated archaeological sediment blocks at a microstratigraphic scale. First, we evaluate the effects of resin impregnation on proteins in skeletal material by encasing modern bone material in resin. We find minimal effects of resin impregnation, with a similar amount of proteomic information, as represented by SPIN site counts, being recovered before and after resin impregnation. We observe an increase in oxidation-related post-translational modifications. Encouraged by these results, we extract proteins from resin-impregnated sediment blocks from the Palaeolithic sites of Bacho Kiro Cave, La Ferrassie and La Grande Roche de la Plématrie at Quinçay (henceforth, Quinçay). We find that, for Bacho Kiro Cave and La Ferrassie, taxonomic identification through SPIN is possible for all samples where bone fragments were targeted, with the identifications corresponding to previous analyses of the faunal community at these sites (12–14). For Quinçay, only two out of eleven bone-derived samples could be identified, likely due to poor preservation of the sampled sedimentary deposits. These results open the detailed study of faunal community composition at the scale of past human occupations through palaeoproteomic analysis of undisturbed archaeological deposits encased in resin-embedded sediment blocks.

## Results

### Effect of Resin Impregnation on Protein Recovery and Modifications

To test the effects of resin impregnation on bone protein extraction, we sampled three modern bones: a *Bos taurus* tibia (specimen 1), a calcaneus, also from *Bos taurus* (specimen 2), and a *Canis lupus* ulna (specimen 3; Figure S6). The bones were first sectioned to expose their interior with a microdrill. Subsequently, using a sterile dentistry drill on a bleach-cleaned surface, we took three powder samples per bone from different locations on the bone’s interior sections (Figure S7). After sampling, the bones were impregnated with a mixture of non-accelerated polyester resin, mixed with styrene and a catalyzer (Methyl Ethyl Ketone Peroxide, MEKP) in a 7:3:1 proportion, respectively. After drying for several days at ~40 ºC, the resin-hardened bones were then re-sampled (three samples per bone), with sample locations adjacent to those taken before resin-impregnation.

The recovered proteomes from the resin experiment were compared between samples taken before and after resin impregnation to evaluate what effects resin impregnation may have on bone proteome recovery. The percentage of identified MS2 spectra varies between 2.9% and 5.4% per sample (Figure 1a), which is a level often seen for archaeological specimens (15). The percentage of identified spectra does not differ depending on whether samples had been resin-impregnated or not, nor between specimens (LM, null model best fit), indicating that resin impregnation does not affect our ability to identify which peptides are present in a sample. The site count based on SPIN (i.e., the number of amino acid positions called in relation to the top-ranking taxonomic identities) varies between 2,693 and 4,856 per sample (Table 1) but does not significantly depend on resin impregnation or specimen either (LM, null model best fit). Blanks from the resin impregnation process have a site count of 758 (before resin-impregnation) and 782 (after resin-impregnation), whereas the protein extraction blank has a site count of 291, indicating a low amount of protein contamination.

**Table 1.** SPIN site counts before and after resin impregnation. Means are given ± 1 standard deviation, based on 3 replicates for each specimen. See photos of specimens in Figure S6.

| Specimen | Taxon (bone element) | Site count before (total) | Site count after (total) | Site count before (NCPs) | Site count after (NCPs) |
| --- | --- | --- | --- | --- | --- |
| #1 | *Canis lupus* (Ulna) | 3,863.0 ± 152.6 | 3,933.0 ± 123.8 | 756.7 ± 86.1 | 715.0 ± 32.4 |
| #2 | *Bos taurus* (Tibia) | 3,833.3 ± 29.7 | 3,590.3 ± 1,127.6 | 682.0 ± 24.2 | 567.7 ± 462.9 |
| #3 | *Bos taurus* (Calcaneus) | 3,700.0 ± 228.8 | 2,963.7 ± 192.4 | 511.3 ± 114.2 | 295.3 ± 136.6 |

**Figure 1.**
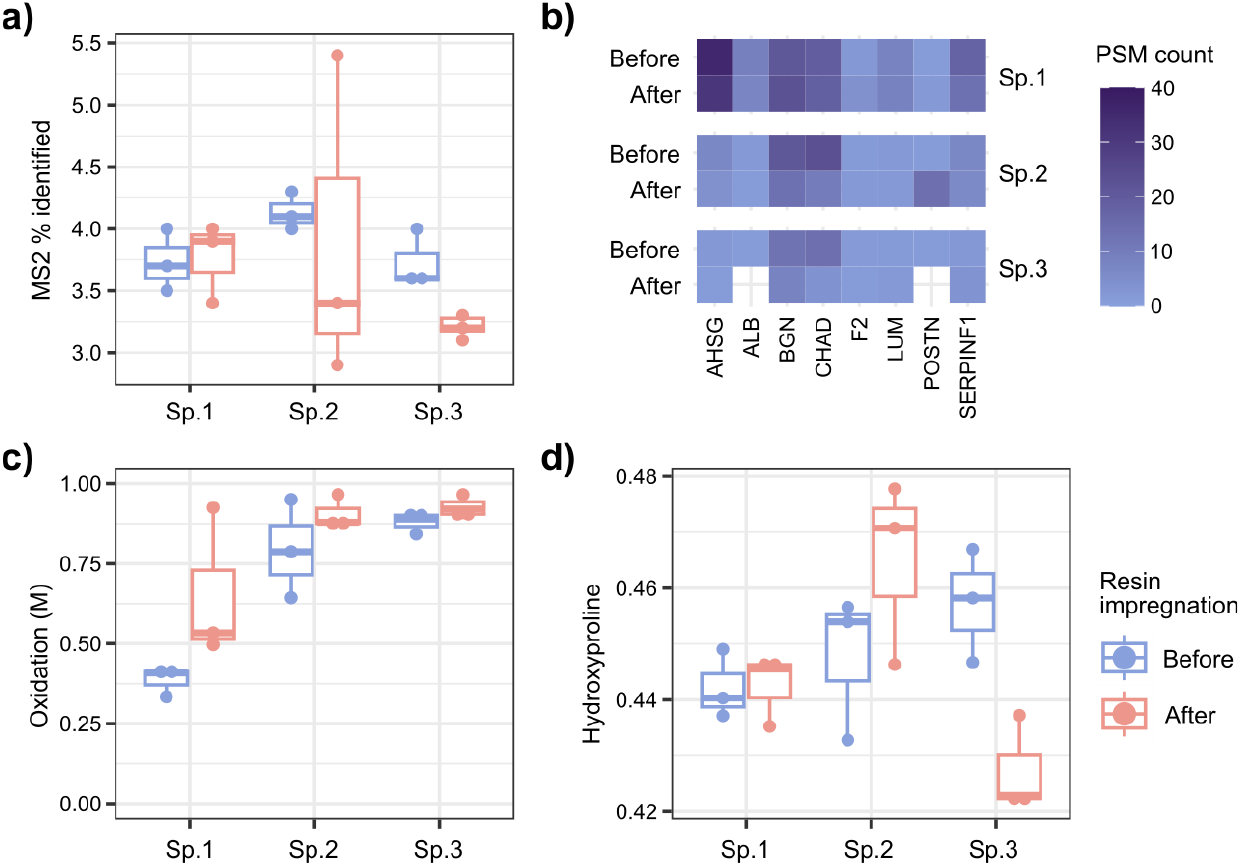
Effects of resin impregnation on proteome recovery and modification. a) The percentage of identified MS2 spectra. b) Summed count of peptide spectrum matches (PSMs) for all identified non-collagenous proteins (NCPs). c) Fraction of oxidated methionines (M). d) Fraction of hydroxylated prolines (P). Sp. = specimen.

For all samples, between 18 and 20 proteins were identified (out of a maximum of 20 proteins included in the SPIN database), but protein number does not vary by resin impregnation or specimen (LM, null model best fit). For the samples where we did not recover all 20 proteins, two were missing ALB (Specimen 2 after; Specimen 3 after), one CHAD (Specimen 3 after), one COL12A1 (Specimen 3 after), and 12 were missing POSTN (five before-samples, seven after-samples). For most NCPs (non-collagenous proteins), fewer PSMs (peptide spectrum matches) were identified after resin impregnation (Figure 1b). This indicates some degree of information loss of less abundant proteins through resin impregnation. Between six and eight NCPs are found in all samples, but the number of NCPs is not affected by resin impregnation (LM, null model best fit). The site count of NCPs only depends on specimen, not resin impregnation (LM, F=4.27 and p=0.034); however, it should be noted that there is a large variability in the NCP site counts between samples, especially within Specimen 2 (Table 1).

In addition to protein and peptide identification, we explored how the proteins are affected by resin impregnation in terms of post-translational modifications (PTMs) and peptide length. The extent of oxidation of methionine (M) (weighted by peptide intensity, calculated for samples with more than 10 peptides with M) depends on both resin impregnation and specimen, and increases after resin impregnation (LM, F=5.21 and p=0.039; Figure 1c). This pattern holds true also when looking at only COL1 (LM, F=6.908 and p=0.020), as well as when restricted to NCPs, although only a few samples have >10 methionine-containing NCP peptides (n=5 samples; Figure S8a-b). For hydroxyproline, as for methionine oxidation, both specimen and resin impregnation have a significant effect, and a significant interaction was found between specimen and resin impregnation (LM, F=7.51 and p=0.008; Figure 1d), with two out of three specimens showing higher levels of hydroxyproline after resin impregnation, and one markedly lower. However, when filtering for only COL1 peptides, this pattern is no longer significant (LM, null model best fit), and the fraction of hydroxylated prolines in NCPs only significantly differs by specimen (LM, F=8.509 and p=0.008; Figure S8c-d). This indicates that the resin impregnation process is increasing oxidation levels of amino acids, especially for methionines. Although there is also an effect of resin impregnation on hydroxyproline levels, the effect is small, in the range of ± 3%. Deamidation of both asparagine (N) and glutamine (Q) (excluding contaminants; Figure S9) differs significantly between specimens, but is not dependent on resin impregnation (LM, F=26.15 and p<0.001 for N; F=28.49 and p<0.001 for Q). Peptide length, weighted by intensity, does not vary depending on resin impregnation or specimen (LM, null model best fit). It therefore appears that sufficient proteome recovery for taxonomic identifications is possible after resin impregnation of bone specimens, even though there is a potential loss of peptides from low-abundance NCPs and an increase in oxidative PTMs.

### Protein Preservation in Pleistocene Archaeological Blocks

As no major negative impact on obtaining skeletal proteomes sufficient for taxonomic identification could be identified from the process of resin impregnation, proteins were extracted from bone fragments embedded in resin-impregnated sediment blocks from three Palaeolithic archaeological sites: Bacho Kiro Cave (Bulgaria), La Ferrassie (France) and Quinçay (France) (Table S1 and Figure S1). In addition to bone fragments, we also sampled the sediment matrix in order to understand skeletal protein preservation and their presence at a microscale.

For the sediment block coming from Bacho Kiro Cave, taxonomic identification through SPIN was possible for all 11 samples that included visually identified bone fragments (see infrared spectra in SI, Figures S11-12). Of these, five were identified as *Bos* sp./*Bison* sp., two as *Equus* sp., two as *Ursus* sp, one as *Ursus arctos* and one as Caprinae (Figure 2). This is in agreement with the taxa previously identified as dominant at the site through ZooMS and zooarchaeological analysis (12, 14). At La Ferrassie, taxonomic identification was also possible for all samples containing visible bone material (Figures S13-14), as 10 out of 11 samples were identified as *Bos* sp./*Bison* sp. and the last sample as Cervidae. Interestingly, taxonomic interpretations were achieved for sample 42 (Figure 3), which is a bone burned to temperatures above ~700 ºC as shown by the presence of the phosphate high temperature peak associated with the hydroxyl (OH) libration band at ca. 630 cm^-1^ (Figure S14; (16), though it should be noted that some sediment adjacent to this bone was also sampled. For La Ferrassie, previous ZooMS analysis has shown a dominance of *Bos* sp./*Bison* sp. and *Rangifer* sp. in the faunal assemblage (12). Since the SPIN database only contains a single Cervidae species, it is likely that the Cervidae assignment corresponds to a *Rangifer* sp. specimen. Finally, out of the 11 samples from Quinçay containing visible bone, only two could be taxonomically identified through SPIN, one sample as *Bos* sp./*Bison* sp. and one as *Bubalus bubalis*/Cervidae (Figure S5). Previous ZooMS analysis of bone fragments from this site has shown a dominance of *Bos* sp./*Bison* sp., Equidae and *Rangifer* sp., which is compatible with the two SPIN identifications (10). Palaeoproteomic analysis of bone specimens from Palaeolithic resin-impregnated sediment blocks thereby has a high success rate at two of the three analysed sites, with good agreement with previous faunal composition analysis through ZooMS and zooarchaeology.

**Figure 2.**
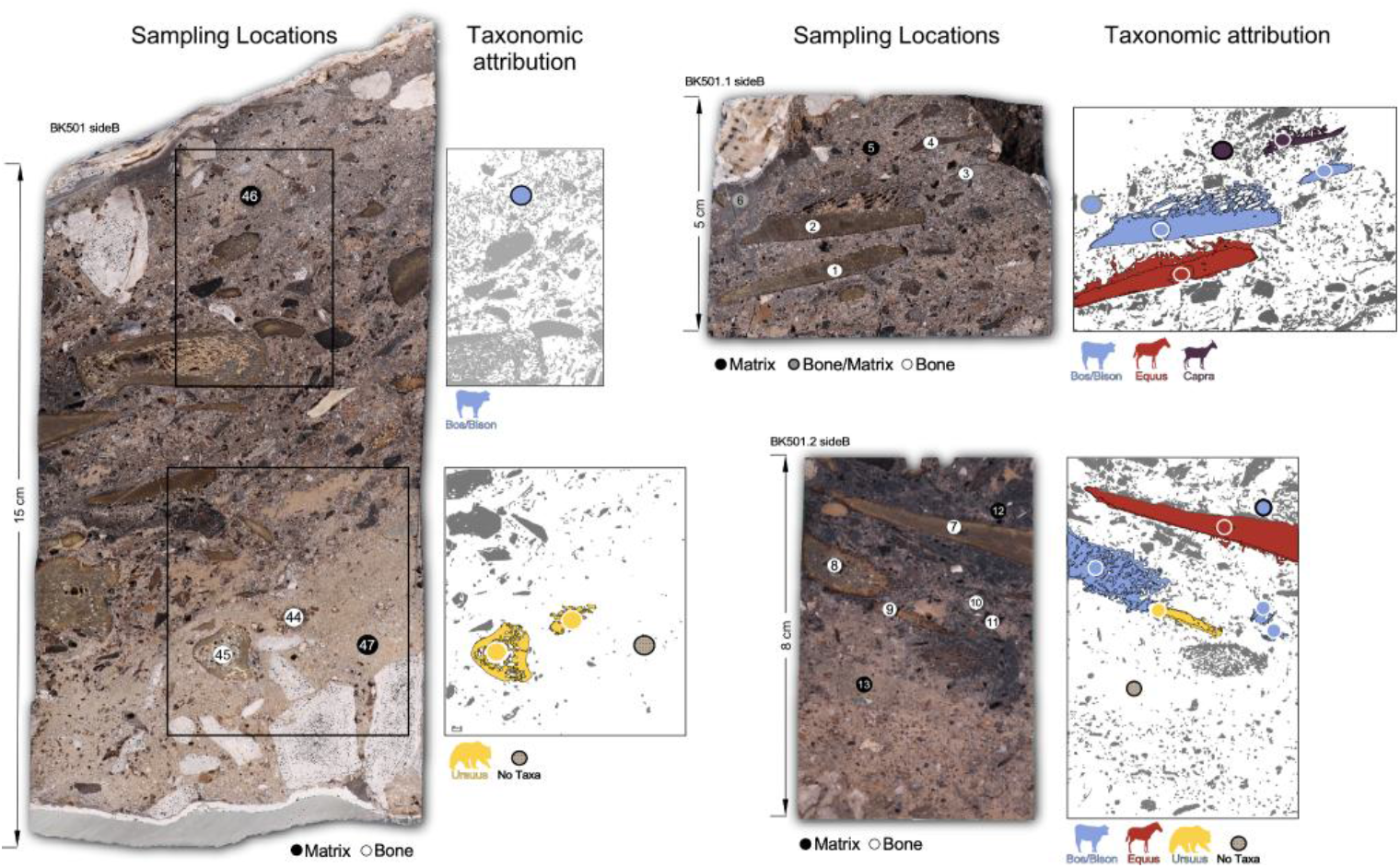
Bacho Kiro Cave sampling locations and taxonomic attributions of block BK 501. Bones in grey were not sampled and bones in colour have specific taxonomic attributions. Several cuts from the same block are indicated. Circles in sampling locations depict the drilled areas and unique sample numbers (see SI Table S1).

**Figure 3.**
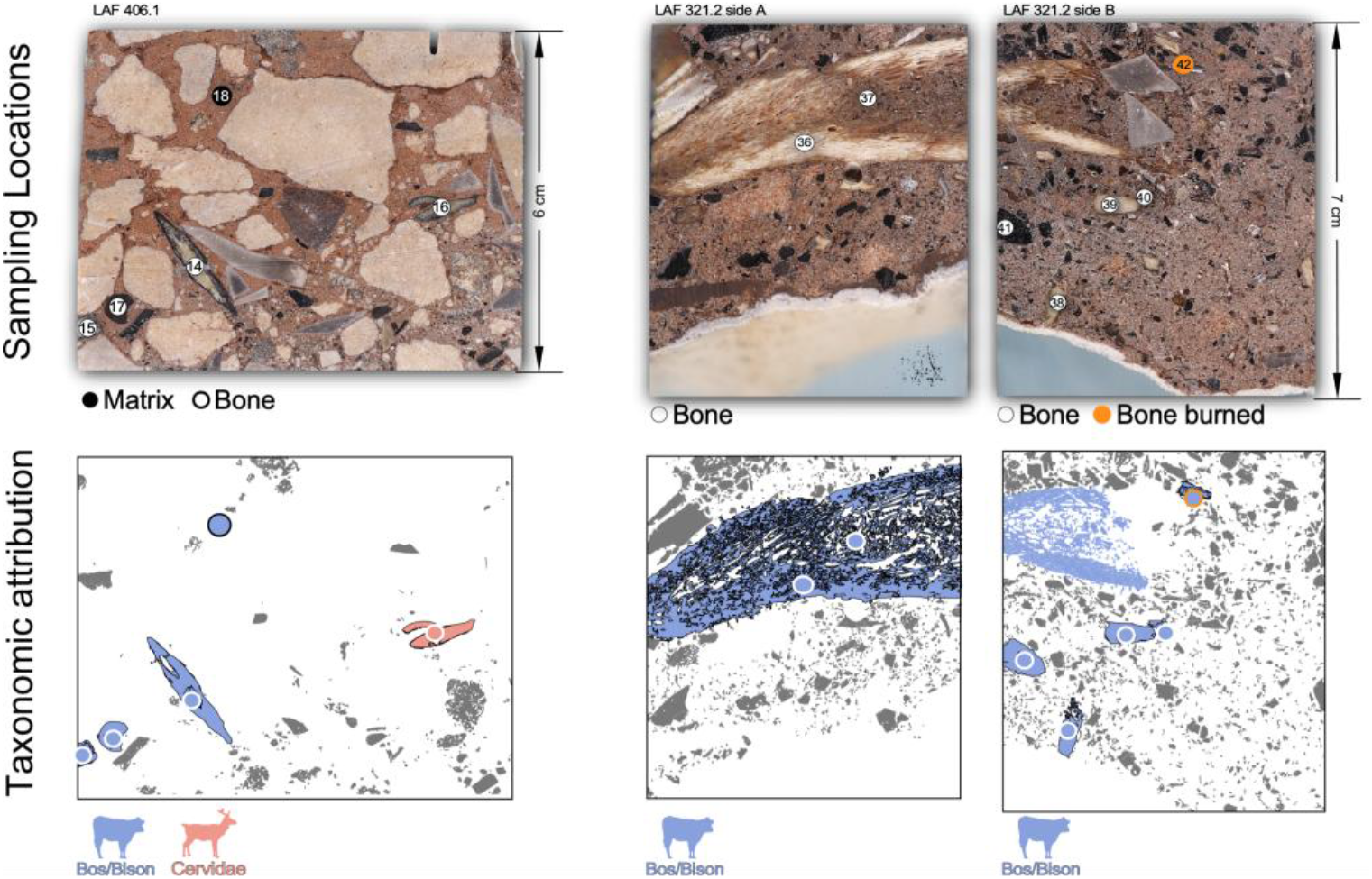
La Ferrassie sampling locations and taxonomic attributions of blocks LAF 406.1 and LAF 321.2. Bones in grey were not sampled and bones in colour have specific taxonomic attributions. *Top:* La Ferrassie sampling locations on resin-impregnated block LAF 406.1 and both faces of block LAF 321.2 (side a and b). Circles in sampling locations depict the drilled areas and unique sample numbers (see SI Table S1). *Bottom:* taxonomic attributions.

To identify materials within the sediment blocks with higher proteomic information content and to trace where the identified proteins are likely originating from, we compared palaeoproteomic results between different samples within the archaeological sites using a range of approaches. Bacho Kiro Cave and La Ferrassie show that, overall, there is a higher SPIN site count from samples with bone than samples with matrix (Figure 4b). For Bacho Kiro Cave, the lowest number of amino acid sites were for black bone (947.0 ± 130.1) and the highest for samples with mixed bone and matrix (3,052.0 ± 553.0). For La Ferrassie, the lowest number of amino acid sites was obtained from the matrix sample (295), whereas the highest number was from pure bone samples (1,780.4 ± 338.0). Sample 42, which was found to be burned to above ~700 ºC (Figures S3 and S14), has the lowest site count of all bone specimens from La Ferrassie (970). It should be noted, however, that this sample was also the smallest (1.0 mg, whereas other bone samples were 4.0-12.6 mg), although no significant relationship between weight and site count was found. Finally, from Quinçay the highest number of amino acid sites was obtained from the black bone sample (468) and the lowest from the pure resin sample (70). These results indicate that the largest amount of proteomic information is preserved in visually identifiable bone fragments, and the taxonomic identifications of bone fragments are unlikely to be significantly influenced by proteins from the surrounding matrix, given their much lower abundance. From the laboratory blank associated with Bacho Kiro Cave, 766 sites were covered, whereas 291 sites were covered in the blanks associated with La Ferrassie and Quinçay. The blanks are therefore on a comparable level to the most data-poor samples from each site.

**Figure 4.**
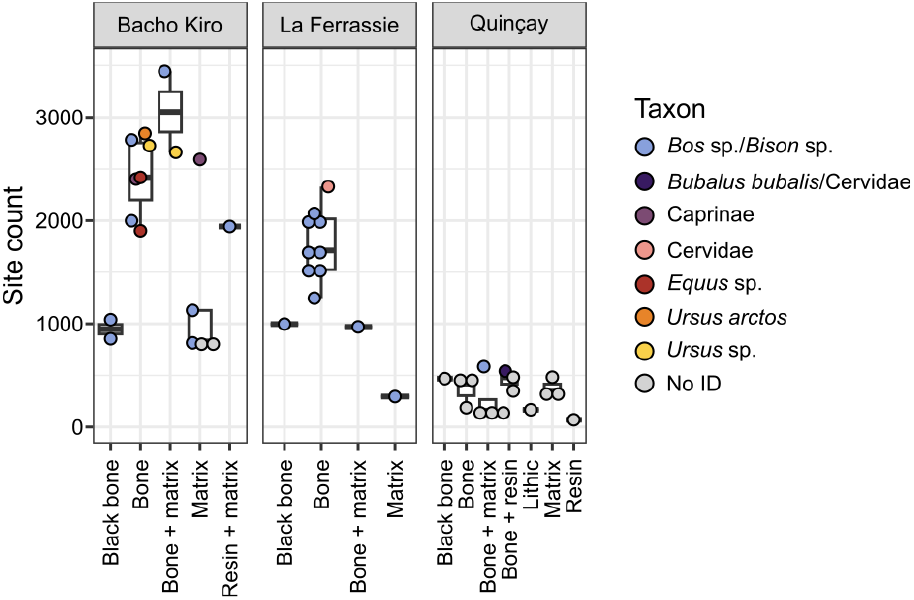
Proteomic information recovery from archaeological sediment blocks. Proteomic information content is represented by the SPIN site count, from each site and sample type. All samples are coloured by the taxonomic identification obtained through SPIN.

It was not possible to determine the taxonomic identity for two samples from Bacho Kiro Cave, both stemming from samples drilled from the matrix of the blocks. Three matrix samples from Bacho Kiro Cave did, however, gain a taxonomic identification through SPIN, with two identified as *Bos* sp./*Bison* sp. (815 and 1,129 sites covered, respectively) and one as Caprinae (2,592 sites). The large number of sites covered in the Caprinae matrix sample, similar to the bone samples from Bacho Kiro Cave, may indicate that a minor amount of bone material was included in this matrix sample, although it was not visually identified. This is further supported by the identification of 11 different collagen proteins in this sample, with a total of 656 PSMs. In contrast, the two matrix samples identified as *Bos* sp./*Bison* sp. have 6 and 7 collagens with 109 and 157 PSMs, respectively. The ubiquitous presence of microscopic-sized bones in the sedimentary matrix is further shown by both micromorphological and μXRF (micro X-ray fluorescence) data exposing microparticles rich in calcium (Ca) and phosphorous (P) associated with the microscopic bones throughout the Bacho Kiro Cave sampled block sediments (Figure S4). A matrix sample from La Ferrassie was identified as *Bos* sp./*Bison* sp. (295 sites) and the laboratory blank associated with La Ferrassie as *Canis lupus* (291 sites). Peptides identified in the matrix-samples stem from various collagens (mainly COL1A1 and COL1A2) but are often taxonomically unspecific. The proteomic content of the matrix samples therefore likely originates from microscopic bone particles.

Deamidation of both asparagine (N) and glutamine (Q) was significantly different both between the archaeological sites of Bacho Kiro Cave and La Ferrassie and between sample types, without a significant interaction between the variables (LM, F=6.24, p=0.005 and F=2.65 and p=0.026, respectively). However, there is no clear trend as to which material type has the highest or lowest deamidation across sites (Figure 5a-b). As deamidation is correlated with protein degradation, this would indicate that no material types contain proteins with enhanced proteomic degradation. For oxidation of methionine and hydroxyproline, no clear patterns are apparent across sites and tissue types (Figure S10a-b). Mean peptide length, weighted by intensity, appears to be highest in samples deriving from bone fragments in the sediment blocks (Figure 5c). This indicates better preservation of proteins within visibly identifiable bone fragments, as also indicated by the higher site counts of such samples (Figure 4).

**Figure 5.**
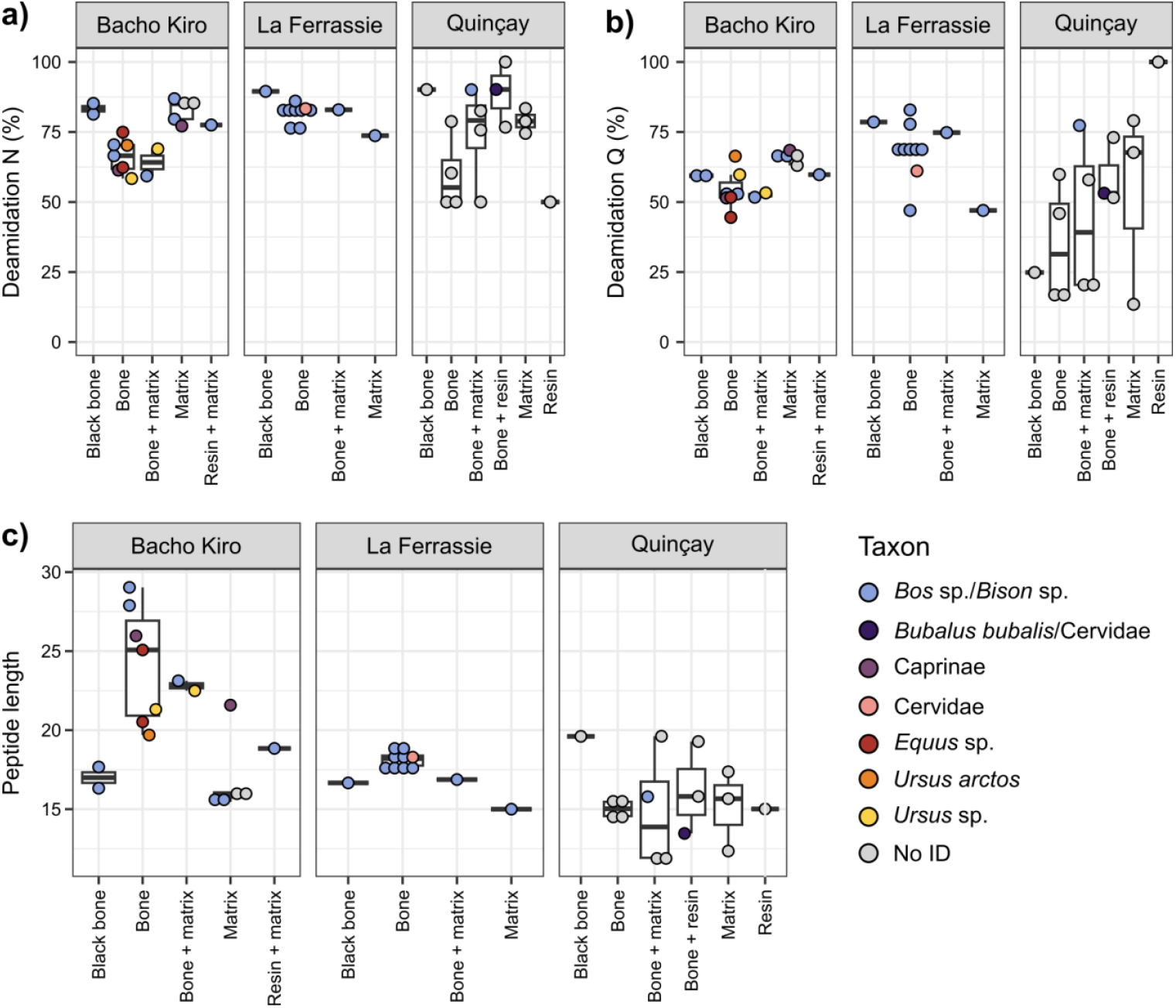
Properties of peptides from archaeological blocks. a) Percentage of deamidated aspargine (N), b) Percentage of deamidated glutamine (Q), and c) Mean peptide length, weighted by peptide intensity. Each point is a sample, colored by taxonomic identification through SPIN.

## Discussion

Advancements in obtaining biomolecular information from archaeological sediments has been one of the most outstanding achievements in archaeological research in the last two decades. However, the microstratigraphic complexity and the heterogeneous nature of archaeological deposits is often not considered in these applications. Analyses tend to be performed at a resolution corresponding to entire stratigraphic layers. Yet, stratigraphic layers often accumulate over hundreds or even thousands of years, which is ample time for several – and potentially distinct – types of human, and faunal, occupations to occur. This palimpsest nature of the archaeological record is well known, but it is not easy to tease apart individual depositional events. The issue of scale is especially acute for periods of transition in human behaviour that are, geologically speaking, not very long. This means that stratigraphic deposits associated with potentially distinct occupations tend to be thin and that distinct occupations can only be studied at a microstratigraphic scale. By using a microarchaeological approach applied to bone fragments embedded in intact resin-impregnated sediment blocks, we can retain this microstratigraphic resolution of the original context, the fine temporal relationship of the deposits, and their postdepositional alterations. Here, we show that palaeoproteomic analysis can be conducted directly on such bone fragments embedded within resin-impregnated sediment blocks at this microstratigraphic scale. Our approach provides the ability to address temporal and contextual relationships between discrete depositions within a stratigraphic layer and provides taxonomic identification of organic remains directly deriving from human activities.

Through experimental work using modern bones, we find that protein recovery is only mildly affected by resin impregnation, seen as a loss of peptides from low-abundance non-collagenous proteins and an increase in oxidative post-translational modifications, the latter especially for methionine and possibly to a far lesser extent for hydroxyproline. Nevertheless, the possible increase in hydroxyproline rates is of relevance when attempting to study similar resin-impregnated bone specimens through lower-resolution protein mass spectrometry where no fragment ion series are generated, such as with MALDI-ToF MS. Since several of the peptide marker masses used in zooarchaeology by mass spectrometry (ZooMS), which generally utilises MALDI-ToF MS technology, are 15.99 Da distant from each other, that is, equal to the oxidative mass changes caused by resin impregnation, such taxonomic identification approaches potentially return false taxonomic identifications in contexts where resin impregnation would have led to significantly higher abundances of hydroxyproline.

From archaeological sediment blocks, taxonomic identification of bone material is highly successful in two out of three analysed Palaeolithic sites, whereas the third block from Quinçay shows poor protein preservation overall. The reasons for this poor preservation can be linked to diagenesis. The area from which this block was sampled is in a natural depression inside the cave, which enhanced water mobility throughout the deposits in combination with guano-rich solutions. As water content, pH and soil chemistry affect protein preservation, this may have been detrimental for protein preservation (17, 18). Additional samples from other locations within Quinçay are needed to test if localized post-depositional processes affected protein preservation differently across the site, as suggested based on protein deamidation from previously ZooMS-analysed bone specimens from Quinçay (13).

Overall, our study shows that even mm-sized bones are excellent sources of proteins in Palaeolithic deposits. Some of the sediment matrices have also provided taxonomic identifications largely based on collagen type I, indicating that microscopic-sized bones in these deposits are ubiquitous and likely responsible for the recovered proteomic data. Some of the bones that provided taxonomic data had black and white colorations – features often associated with charring and calcination, respectively (19, 20). However, further work is needed to test the preservation of proteins in samples affected by burning.

Our results show distinct patterns of taxon distribution across the deposits of Bacho Kiro Cave and La Ferrassie. At Bacho Kiro Cave, a higher diversity of taxa (n=4) was identified, with bones retrieved from specimens in close spatial relationships (just a few mm and cm apart). The diverse taxonomic attributions support the jumbled nature of materials resulting from debris derived from human actions (such as dumping) in Layer N1-I (21). At La Ferrassie, however, only one sample was assigned to a Cervidae, with all the other sampled bones pertaining to the same taxon (*Bos* sp./*Bison* sp.), even though two distinct blocks from layers several meters apart were sampled (Figure S2). Our results are consistent with previous ZooMS results for Layer 6 of La Ferrassie, showing that *Bos* sp./*Bison* sp. are the most abundant taxa from the unidentifiable, fragmented fauna, despite *Bos* sp./*Bison* sp. not being the main taxa determined morphologically (12). It should be noted, therefore, that although the palaeoproteomic analysis of bone fragments at a microstratigraphic scale opens various new areas of research, the incorporation of this information into quantitative zooarchaeological frameworks is likely to be complicated, as recently encountered when integrating ZooMS and zooarchaeological NISP counts (7, 22).

The ability to extract proteins from resin-impregnated sediments opens new research avenues and there are research questions for which we believe this microscale resolution would be the best approach. Of these, we highlight here four main areas of future research.

1. Obtaining microstratigraphic resolution of distinct occupations – As mentioned above, the characterization of faunal assemblages is typically performed using a stratigraphic layer as the analytical unit. However, these associations have unknown degrees of micro-contextual accuracy. We all acknowledge that, within a stratigraphic layer, there are several, potentially very distinct, superimposed occupations that are impossible to separate during excavations. Considering the microstratigraphic complexity of archaeological sites, extracting proteins from multiple resin-encased deposits allows for proveniencing and identifying bones with a microstratigraphic resolution. The resulting disentanglement of time-distinct depositions is particularly relevant for interpreting discrete site use changes through time and identifying behavioral activities (e.g., species-specific processing areas, accumulation of occupation detritus over time, hearth-related accumulations) within a layer. The ability to reveal the faunal composition of ‘micro-assemblages’ within a resolvable layer should reflect upon source and mode of deposition of these elements, a subtle aspect of site formation processes.
2. Determining the context of hominin fossils – If the research goal is to find, often elusive, human remains from unidentified bone fragments, then focusing on bone fragments placed within resin-impregnated blocks may not be an advantageous approach, as these blocks only cover a spatially limited area within a site. Notwithstanding, we would argue that if a hominin bone were to be identified in resin-impregnated sediments, the degree into which we can investigate the depositional context of that fossil is unprecedented. The resin-encased hominin bone would still be preserved in its original depositional context, allowing for a better understanding of its microstratigraphic environment and associated site formation processes. For instance, claims for intentional burials are often heatedly disputed (e.g., (23–27)) and having fine-scale microstratigraphy associated with a hominin fossil is essential to access its context.
3. Targeting microscopic-sized bones – The sample size used for the outlined method is very small (up to 10 mg), with the positive taxonomic results from the matrix samples suggesting that microscopic-sized bones can be targeted in the future. This is relevant as it permits the identification of bones that normally are not recovered even in the screened fractions of archaeological excavations. Identifying small-sized bone fragments might be relevant in order to investigate fragmentation-level discrepancies when compared with the macrofauna, as for instance with the degree of fragmentation derived from anthropic actions (e.g., bone splintering for grease rendering, intensity of trampling in a given area). While infrequent, we do have archaeological sites where small, fragmented bones are still embedded in the sediments although macroscopic bones are poorly or not preserved. Targeting those small bone fragments can then provide an otherwise non-existent zooarchaeological record.
4. Assessing taphonomic changes – Extracting proteins from undisturbed bones opens the possibility of contextualizing the geochemical depositional context associated with protein preservation – the question of when/how deamidation occurs seems variable, in part depending on the micro-depositional environment and localized geochemical conditions. The nature and geochemical signatures of sediments embedding bones can be assessed from extracting proteins from bones in their intact context in the resin-impregnated blocks, particularly when protein extraction is used in tandem with non-destructive techniques such as micro-FTIR for mineralogical characterization or μXRF for elemental mapping.

By studying a small number of Late Pleistocene, well-contextualised micromorphology blocks, and supported by FTIR and μXRF, we demonstrate that tandem mass spectrometry is capable of providing taxonomic identifications of bone fragments embedded in resin-impregnated micromorphology blocks. The obtained taxonomic identifications, sometimes of bone fragments deposited millimeters apart from each other, are in general agreement with previous proteomic and zooarchaeological research at the three sites studied. Our results also showcase that bone particles that are invisible to the bare eye, embedded in the sediment matrix, can likewise provide taxonomic hints of past hominin behaviour and/or fauna community composition at Late Pleistocene sites. Finally, the combination of FTIR, μXRF, and LC-MS/MS hints at the possibility to study through mass spectrometry Late Pleistocene bone fragments that have been heated. Together, this paves the way for a range of future palaeoproteomic applications at a resolution that matches the time scales at which past human occupations occurred.

## Materials and Methods

### Impregnation methodology

Samples used to extract proteins were collected from archaeological deposits as intact blocks. This technique is typically used for micromorphological analysis, allowing for the preservation of the original association and geometry of the archaeological and sedimentary components (Courty et al., 1989; Goldberg & Aldeias, 2018). Sediment blocks were carved from excavation areas or profiles and extracted as undisturbed volumes, either by wrapping the sediments in soft paper or by using pre-plastered bandages to ensure sample stability. The samples were then dried at ~40ºC for several days and subsequently hardened using a mixture of non-promoted polyester resin diluted with styrene to which a catalyst (MEKP) was added. The hardened samples were then cut and trimmed into several slabs, some of which were used to produce thin sections for soil micromorphological analysis by Spectrum Petrographics Inc. (Vancouver, Washington) and by Thomas Beckmann (Germany). For protein extraction, we used the leftover resin-impregnated slabs, with targeting of several bone or matrix samples within each slab. The remaining hardened slabs were saved and stored at the ArchaeoDirt laboratory at the International Center for Archaeology and the Evolution of Human Behavior (ICArEHB) at University of Algarve (Portugal).

### Archaeological case-studies

We selected resin-hardened slabs from three different Pleistocene cave sites for protein extraction, namely Bacho Kiro Cave (Bulgaria), La Ferrassie (France), and Quinçay (France). Table S1 presents the list of samples used in this study.

#### Bacho Kiro Cave

Bacho Kiro Cave is located near the town of Dryanovo in Bulgaria on the northern range of the central Balkan Mountains. The site was initially excavated during the twentieth century, first by D. Garrod and R. Popov in the 1930s and in the 1970’s by a Bulgarian-Polish team led by B. Ginter and J. Kozłowski (28, 29). More recent excavations (2015-2021) were conducted by a joint effort between the National Archaeological Institute with the Museum of the Bulgarian Academy of Sciences (Sofia, Bulgaria) and the Department of Human Evolution at the Max Planck Institute for Evolutionary Anthropology (Leipzig, Germany) (21). Archaeological remains are present near the entrance of a larger karstic system formed on Cretaceous sandy limestones that form a canyon along the Dryanovo River. New excavations focused on two sectors, the Niche 1 and the Main sector (21). The stratigraphic sequence starts at the bottom with Middle Paleolithic occupations (Layer K) dated to 61 ± 6 ka (OSL) (30) and to >51 ka cal BP (31). These deposits are subsequently overlain by occupations associated with techno-cultural artefact assemblages identified as a variant of the Initial Upper Paleolithic found at the top of Layer J and within Layer I. These assemblages are dated to ~45-42 ka cal BP (31, 32). Archaeological remains above these layers are sparse but attributable to Aurignacian industries, dated to 36-34 ka cal BP (31). At the top, the sequence ends with layers attributable to Epigravettian.

The resin-impregnated samples used in this study correspond to samples collected from the excavation area of the Niche 1 and spanning the Layers I and J. Zooarchaeological studies on identified bone from layers I and J showed the predominance of *Bos* sp./*Bison* sp., cave bear (*Ursus spelaeus*) and red deer (*Cervus elaphus*) with minor representation of additional carnivores and herbivores (14).

#### La Ferrassie

The site of La Ferrassie is located along a tributary valley to the Vézère River near the town of Les Eyzies-de-Tayac in the Dordogne area of southwestern France. The site is carved into Upper Cretaceous carbonates that lie unconformably on Upper Jurassic limestones. La Ferrassie has a long history of research, starting with limited (and unpublished) work in the late 19^th^ century, followed by extensive excavations by Capitan and Peyrony in the beginning of the twentieth century (33, 34), and then by excavations by H. Delporte in the 1970s (35). More recent excavations directed by A. Turq and colleagues from 2010 to 2014 focused on two areas, the Western Sector and the Northern Sector (36). La Ferrassie is well-known for the discovery of several Neanderthal individuals as well as for its long sequence of both Middle and Upper Paleolithic occupations. Recent geoarchaeological studies have shown that the so-called “grand abri” of La Ferrassie was, at its origin, a large cave with its entrance in the area of the Western Sector (37). The stratigraphic sequence of the Western Sector is subdivided into nine major stratigraphic units. The basal occupations are Middle Paleolithic and span MIS5 (Layer 1), MIS4 (Layer 2), early to mid MIS3 (Layers 3, 4, 5a and 5b) - (38, 39). The Middle Paleolithic deposits are then capped by Châtelperronian assemblages (Layer 6) dated to between 45,100 and 39,520 cal BP (39) and overlaid by Aurignacian occupations (Layers 7a and 7b).

The resin-hardened blocks selected for this study are from the uppermost Mousterian Layer 5b, characterized by a Mousterian with a relatively high Levallois component. These sediments contain abundant anthropogenic inputs, namely burned and unburned bone fragments. The faunal spectra of the main prey taxa are dominated by auroch/bison (*Bos* sp./*Bison* sp.) and red deer (*Cervus elaphus*) with minor proportions of reindeer (*Rangifer tarandus*), roe deer (*Capraeolus capraelus*) and horse (*Equus*) throughout all the Middle Paleolithic layers (12, 40).

#### La Grande-Roche-de-la-Plématrie (Quinçay)

The site of Quinçay – also known as La Grande Roche de La Plématrie (GRL) – is located 10 km west of the town of Poitiers (Vienne, France). The site is a small limestone cave (~20 m deep x 10 m wide chamber) facing a dry valley that is a tributary of the Auxance River. Initially discovered in 1952, the site was excavated between 1968 to 1990 by F. Lévèque (41, 42) and is currently the target of renewed excavations by M. Soressi. More than 18 meters of profiles are exposed, albeit not always preserving the totality of the circa 1,5 meter high sequence, since the lower archaeological layers near the cave entrance are physically disconnected from those towards the back of the cave due to the rising of the bedrock. The sequence contains Mousterian assemblages (43) in the lower layers with little or no bone preservation, followed by Châtelperronian assemblages associated with abundant lithics and bone fragments, including bone tools (44) and ornaments (45). As part of ongoing work at the site, the stratigraphic sequence and associated technocomplexes at Quinçay are currently reassessed. Previous work on ZooMS showed the dominance of horse (*Equus*), auroch/bison (*Bos* sp./*Bison* sp.) and reindeer (*Rangifer*) (13).

The selected resin-hardened blocks from Quinçay were collected from the frontal profile (FRON1) towards the back of the cave from Châtelperronian layer 30b and the underlying archaeologically less rich layer 40. No bone specimens from the depth at which the micromorphology block was taken were studied by ZooMS previously (13) Recent excavations recovered several bone specimens in the immediate vicinity of the micromorphology block studied here.

### Protein extraction

Laboratory work was conducted in facilities dedicated to the processing of ancient biomolecules at the Globe Institute, University of Copenhagen (Denmark). Micromorphology block surfaces were briefly rinsed with diluted bleach, neutralised with water, then air-dried. Extraction blanks were carried alongside the samples, one per extraction batch, in order to monitor laboratory contamination. Proteins were extracted following a modified AmBic-acid extraction protocol (10, 46). Briefly, approximately 10 mg of bone/resin powder was subsampled whenever possible (Supplementary Data 1). The samples were suspended in 100 μl of 50 mM ammonium bicarbonate (AmBic) and incubated at 65 °C for 1 h. The supernatant was thereafter collected, and 100 μl of 5% hydrochloric acid (HCl) was added to the sample pellet. The samples were demineralized at 4 °C overnight, whereafter the supernatant was removed. The remaining acid-insoluble fraction was washed three times with 100 μl of 50 mM AmBic, in order to remove any remaining acid. A new 100 μl of 50 mM AmBic was added to the samples, and gelatinized at 65 °C for 1 h. Thereafter, 50 μl of AmBic supernatant was combined with 50 μl of the AmBic supernatant which was previously collected. Digestion was carried out overnight using 0.8 μg of sequencing grade modified trypsin (Promega) at 37 °C and 300 rpm agitation. Digestion was stopped through adding 5 μl of 5% trifluoroacetic acid (TFA). Peptides were cleaned using Evotip C18 filters (Evosep) following the manufacturer’s instructions, using 20 μl of peptide extract per sample for all archaeological samples and 10 μl for all modern samples (resin experiment).

### Mass spectrometry

Mass spectrometry was performed in two different batches. The samples from the Bacho Kiro sediment blocks were analysed at the Centre for Protein Research (University of Copenhagen, Denmark) and the samples from the resin experiment, La Ferrassie, and Quinçay at the Proteomics Research Infrastructure (University of Copenhagen, Denmark). In the case of the data generated at the Centre for Protein Research, data was acquired on an Exploris 480 instrument in data-dependent acquisition mode, coupled to an EvoSep nanospray in 100SPD mode. Mass spectrometry was performed in positive ionisation mode at 2 kV, with the ion transfer tube at 275 °C. Scan range of the MS1 was 350-1,400 m/z at an orbitrap resolution of 60,000. The MS2 experiment had 15,000 orbitrap resolution, with high-energy collision dissociation (HCD) and a normalised collision energy of 30% and an isolation width of 1.3 m/z. Top 12 ions with intensity over 200k and charge state 2-6 were fragmented, and thereafter excluded for 30s. In the case of the latter, data was acquired on a timsTOF instrument in data-dependent acquisition mode, coupled to an EvoSep nanospray in 30SPD mode. In the case of the data generated at the Proteomics Research Infrastructure, peptides were separated on a 15 cm × 150 μm ID Pepsep column packed with 1.5 μm C18 beads using the Evosep One HPLC system with the default 30-SPD (30 samples per day) method. The column temperature was maintained at 50 °C. Eluted peptides were ionized via a CaptiveSpray source with a 20 μm emitter and analyzed on a timsTOF Pro2 mass spectrometer (Bruker) operated in PASEF mode. MS data were acquired over an m/z range of 100–1,700 with a TIMS mobility range of 0.6–1.6 1/K_0_. Ion mobility calibration was performed using three Agilent ESI-L Tuning Mix ions: 622.0289, 922.0097, and 1221.9906. The TIMS ramp and accumulation times were each set to 100 ms, with 10 PASEF ramps recorded per 1.17 s cycle. The MS/MS target intensity was set to 20,000, with an intensity threshold of 2,500. A dynamic exclusion of 0.4 min was applied for precursors within a 0.015 m/z and 0.015 V cm^−2^ width window. Given the differences in mass spectrometry approach, data acquisition parameters cannot be compared directly between sample sets, although aggregated and averaged metrics such as site counts and the extent of deamidation should be comparable between samples and specimens. Aside from one sample from Quinçay, mass spectrometry was not performed on peptide extracts from additional pure resin samples, to avoid negative impact on the mass spectrometry equipment.

### Data analysis

Raw data was analysed using MaxQuant v.2.1.3.0 (47). Each archaeological site was searched separately, as was the resin experiment. Specific searches were conducted, using pyro-Glu formation from glutamine and glutamic acid, oxidation (M), hydroxyproline, and deamidation (NQ) as variable modifications. A minimum Andromeda score of 40 was set for modified and unmodified peptides, with FDRs set to 0.01. The data analysis searches, performed separately per experiment and archaeological site, were conducted against the SPIN protein database, containing sequence entries for 20 bone-derived proteins for over 200 mammalian species (6). The MaxQuant internal contaminant database was also included in the search. The mass spectrometry proteomics data, as well as deamidation, MaxQuant and SPIN output, have been deposited to the ProteomeXchange Consortium via the PRIDE (48) partner repository with the dataset identifier PXD061447.

Taxonomic identifications were performed by running the R script provided by Rüther et al. (6), utilising the evidence.txt and summary.txt files from MaxQuant output as the input. SPIN output was analyzed and visualized using R v.4.3.0 (49) using packages *tidyverse* v.2.0.0 (50), *janitor* v.2.2.0 (51), *ggpubr* v.0.6.0 (52) and *metbrewer* v.0.2.0 (53). Deamidation was calculated using the script published in (54). Other PTMs were calculated as a fraction of the respective amino acid that was modified, weighted by peptide intensity. Only samples with more than ten peptides containing the amino acid of interest were included. Statistical tests were conducted using linear mixed effect models with packages *lmerTest* v.3.1.3 (55), *lme4* v.1.1.33 (56) and *MASS* v.7.3.58.4 (57). For the resin experiment, experiment phase (before/after resin impregnation) and specimen were used as fixed effects, and sample weight as a random effect when necessary. For the archaeological blocks, archaeological site and material type were used as fixed effects, and sample weight as random effect when necessary (i.e., if sample weight is not indicated as a random effect, it was not needed to explain the results). Tests including random effects are denoted “LME” and tests without random effects “LM”. Blanks were not included in statistical tests. Code used for the analysis has been archived through Zenodo (10.5281/zenodo.14972505).

### Infrared Spectroscopy

Samples were analysed for FTIR spectroscopy by KBr pressed discs using a Nicolet iS5 spectrometer and OMNIC software (Thermo Fisher Scientific). Approximately 5 mg of bulk sampled drilled combined with KBr was ground to a fine and homogeneous powder using an agate pellet and mortar. Measurements were conducted in transmission mode across the 4,000-400 cm^-1^ range, with a total of 64 scans at a resolution of 4 cm^-1^, with the results reported in absorbance values. The spectra were compared with available databases,with analysis related to heat-induced changes identified based on the presence of an absorbance band at 630 cm^-1^ associated with the hydroxyl groups (OH-).

### μXRF of Block Samples

Elemental mapping of blocks was done using a Bruker M4 Tornado Plus under vacuum under ~2 mbar pressure and using a Rhodium x-ray tube. The acquisition size was 1,700 pixels in width and 2,200 in height. Pixel size was 20 µm with 20 ms per pixel. The elemental maps of calcium, phosphorus and silicon were analysed qualitatively focused on spatial distribution and presence and absence. The produced elemental maps of Ca and P were then used to isolate bone fragments within the resin-impregnated blocks.

## Acknowledgements

We sincerely thank Anna Rufà for generously providing the modern reference bones used in this study. We also thank Pedro Coxito, Miguel Soares Remiseiro, and Carli Peters for their invaluable assistance with the laboratory work, particularly in conducting the μXRF and FTIR analyses. This research is funded by the European Research Council grants MATRIX (project no. 101041245) awarded to V.A., and PROSPER (no. 948365), awarded to F.W., as well as the European Union’s Horizon Europe research and innovation programme under the Marie Skłodowska-Curie grant agreement no. 101106627 (PROMISE, awarded to Z.F.). Views and opinions expressed are however those of the author(s) only and do not necessarily reflect those of the European Union or the European Research Council Executive Agency. Neither the European Union nor the granting authority can be held responsible for them. Mass spectrometry-based proteomics analyses were performed by the Proteomics Research Infrastructure (PRI) and the Novo Nordisk Foundation Center for Protein Research (CPR) at the University of Copenhagen (UCPH), supported by the Novo Nordisk Foundation (NNF) (grant agreement numbers NNF19SA0059305 and NNF14CC0001, respectively). Contribution of M.S. and archaeological work at the site of Quinçay was funded by the Dutch Research council (NWO) ‘Neanderthal Legacy’ grant (VI.C.191.070). Archaeological work at La Ferrassie was funded by the Leakey Foundation, Max Planck Society, Ministère de la Culture (Service Regional de l’Archéologie, Musée Nationale de Préhistoire des Eyzies), Conseil Général de Dordogne, the University of Pennsylvania Museum and the National Science Foundation (Grant #BCS-1755237). The archaeological re-excavation of Bacho Kiro Cave was supported by funding from the Max Planck Society and is a collaborative project between the National Institute of Archaeology and Museum, Bulgarian Academy of Sciences (Sofia) and the Department of Human Evolution at the Max Planck Institute for Evolutionary Anthropology (Leipzig, Germany).

## References

1. M. W. Morley, et al., Why the geosciences are becoming increasingly vital to the interpretation of the human evolutionary record. Nat. Ecol. Evol. 7, 1971–1977 (2023).

2. V. Aldeias, M. C. Stahlschmidt, Sediment DNA can revolutionize archaeology-if it is used the right way. Proc. Natl. Acad. Sci. U. S. A. 121, e2317042121 (2024).

3. C. Rodríguez de Vera, et al., Micro-contextual identification of archaeological lipid biomarkers using resin-impregnated sediment slabs. Sci. Rep. 10, 20574 (2020).

4. D. Massilani, et al., Microstratigraphic preservation of ancient faunal and hominin DNA in Pleistocene cave sediments. Proc. Natl. Acad. Sci. U. S. A. 119, e2113666118 (2022).

5. M. Buckley, M. Collins, J. Thomas-Oates, J. C. Wilson, Species identification by analysis of bone collagen using matrix-assisted laser desorption/ionisation time-of-flight mass spectrometry. Rapid Commun. Mass Spectrom. 23, 3843–3854 (2009).

6. P. L. Rüther, et al., SPIN enables high throughput species identification of archaeological bone by proteomics. Nat. Commun. 13, 2458 (2022).

7. G. M. Smith, K. Ruebens, V. Sinet-Mathiot, F. Welker, Towards a Deeper Integration of ZooMS and Zooarchaeology at Palaeolithic Sites: Current Challenges and Future Directions: Special Issue: Integrating ZooMS and Zooarchaeology. PaleoAnthropology 2024, 186–211 (2024).

8. S. Brown, et al., Identification of a new hominin bone from Denisova Cave, Siberia using collagen fingerprinting and mitochondrial DNA analysis. Sci. Rep. 6, 23559 (2016).

9. H. Xia, et al., Middle and Late Pleistocene Denisovan subsistence at Baishiya Karst Cave. Nature 1–6 (2024).

10. F. Welker, et al., Palaeoproteomic evidence identifies archaic hominins associated with the Châtelperronian at the Grotte du Renne. Proc. Natl. Acad. Sci. U. S. A. 113, 11162–11167 (2016).

11. T. Devièse, et al., Direct dating of Neanderthal remains from the site of Vindija Cave and implications for the Middle to Upper Paleolithic transition. Proc. Natl. Acad. Sci. U. S. A. 114, 10606–10611 (2017).

12. V. Sinet-Mathiot, et al., Identifying the unidentified fauna enhances insights into hominin subsistence strategies during the Middle to Upper Palaeolithic transition. Archaeol. Anthropol. Sci. 15, 139 (2023).

13. F. Welker, M. A. Soressi, M. Roussel, Variations in glutamine deamidation for a Châtelperronian bone assemblage as measured by peptide mass fingerprinting of collagen. STAR: Science & (2017).

14. G. M. Smith, et al., Subsistence behavior during the Initial Upper Paleolithic in Europe: Site use, dietary practice, and carnivore exploitation at Bacho Kiro Cave (Bulgaria). J. Hum. Evol. 161, 103074 (2021).

15. Y. Chiang, F. Welker, M. J. Collins, Spectra without stories: reporting 94% dark and unidentified ancient proteomes. Open Res Eur 4, 71 (2024).

16. C. L. Shaw, An evaluation of the infrared 630 cm−1 OH libration band in bone mineral as evidence of fire in the archaeological record. J. Archaeol. Sci. Rep. 46, 103655 (2022).

17. M. J. Collins, et al., The survival of organic matter in bone: a review. Archaeometry 44, 383–394 (2002).

18. C. Warinner, K. Korzow Richter, M. J. Collins, Paleoproteomics. Chem. Rev. 122, 13401–13446 (2022).

19. M. C. Stiner, S. L. Kuhn, S. Weiner, O. Bar-Yosef, Differential burning, recrystallization, and fragmentation of archaeological bone. J. Archaeol. Sci. 22, 223–237 (1995).

20. G. Gallo, et al., Characterization of structural changes in modern and archaeological burnt bone: Implications for differential preservation bias. PLoS One 16, e0254529 (2021).

21. J.-J. Hublin, et al., Initial Upper Palaeolithic Homo sapiens from Bacho Kiro Cave, Bulgaria. Nature 581, 299–302 (2020).

22. E. Discamps, K. Ruebens, G. Smith, J.-J. Hublin, Can ZooMS help assess species abundance in highly fragmented bone assemblages? Integrating morphological and proteomic identifications for the calculation of an adjusted ZooMS …. PaleoAnthropology (2024).

23. H. L. Dibble, et al., A critical look at evidence from La Chapelle-aux-Saints supporting an intentional Neandertal burial. J. Archaeol. Sci. 53, 649–657 (2015).

24. W. Rendu, et al., Let the dead speak…comments on Dibble et al.’s reply to “Evidence supporting an intentional burial at La Chapelle-aux-Saints.” J. Archaeol. Sci. 69, 12–20 (2016).

25. R. H. Gargett, et al., Grave shortcomings: The evidence for Neandertal burial [and comments and reply]. Curr. Anthropol. 30, 157–190 (1989).

26. D. M. Sandgathe, H. L. Dibble, P. Goldberg, S. P. McPherron, The Roc de Marsal Neandertal child: a reassessment of its status as a deliberate burial. J. Hum. Evol. 61, 243–253 (2011).

27. E. Pomeroy, et al., New Neanderthal remains associated with the “flower burial” at Shanidar Cave. Antiquity 94, 11–26 (2020).

28. D. A. E. Garrod, B. Howe, J. H. Gaul, Excavations in the cave of Bacho Kiro, north-east Bulgaria. Bulletin of the American School of Prehistoric research 15, 46–126 (1939).

29. J. K. Kozlowski, Excavation in the Bacho Kiro Cave (Bulgaria): Final Report. KIP Articles 1893 (1982).

30. S. Pederzani, et al., Subarctic climate for the earliest Homo sapiens in Europe. Sci. Adv. 7, eabi4642 (2021).

31. H. Fewlass, et al., A 14C chronology for the Middle to Upper Palaeolithic transition at Bacho Kiro Cave, Bulgaria. Nat. Ecol. Evol. 4, 794–801 (2020).

32. T. Tsanova, et al., Curated character of the Initial Upper Palaeolithic lithic artefact assemblages in Bacho Kiro Cave (Bulgaria). PLoS One 19, e0307435 (2024).

33. L. Capitan, D. Peyrony, Station préhistorique de la Ferrassie, commune de Savignac-du-Bugue (Dordogne). (1912).

34. D. Peyrony, La Ferrassie. Moustérien, Périgordien, Aurignacien (Editions Leroux, 1934).

35. H. Delporte, “L’Aurignacien de la Ferrassie” in Le Grand Abri de La Ferrassie, H. Delporte, Ed. (Études Quaternaires, 1984), pp. 145–234.

36. A. Turq, et al., Reprise des fouilles dans la partie ouest du gisement de la Ferrassie, Savignac-de-Miremont, Dordogne: problématique et premiers résultats. 78–87 (2012).

37. V. Aldeias, et al., Site formation histories and context of human occupations at the Paleolithic site of La Ferrassie (Dordogne, France). J. Paleolit. Archaeol. 6, 1–63 (2023).

38. G. Guérin, et al., A multi-method luminescence dating of the Palaeolithic sequence of La Ferrassie based on new excavations adjacent to the La Ferrassie 1 and 2 skeletons. J. Archaeol. Sci. 58, 147–166 (2015).

39. S. Talamo, et al., The new 14 C chronology for the Palaeolithic site of La Ferrassie, France: the disappearance of Neanderthals and the arrival of Homo sapiens in France. J. Quat. Sci. 35, 961–973 (2020).

40. S. Pederzani, et al., Reconstructing Late Pleistocene paleoclimate at the scale of human behavior: an example from the Neandertal occupation of La Ferrassie (France). Sci. Rep. 11, 1419 (2021).

41. F. Lévèque, Note à propos de trois gisements Castel-perroniens de Poitou-Charente. Dialektikê. Cahiers de typologie analytique (1980).

42. F. Leveque, J. C. Miskovsky, Le Castelperronien dans son environnement géologique. Essai de synthèse à partir de l’étude lithostratigraphique du remplissage de la grotte de la grande Roche de la Plématrie (Quincay, Vienne) et d’autres dépôts actuellement mis au jour. Anthropologie (L’)(Paris) 87, 369–391 (1983).

43. M. Roussel, M. Soressi, La Grande Roche de la Plématrie à Quinçay (Vienne). L’évolution du Châtelperronien revisitée. Préhistoire entre Vienne et Charente - Hommes et sociétés du Paléolithique 203–219 (2010).

44. M. Roussel, M. Soressi, J.-J. Hublin, The Châtelperronian conundrum: Blade and bladelet lithic technologies from Quinçay, France. J. Hum. Evol. 95, 13–32 (2016).

45. J.-M. Granger, F. Lévêque, Parure castelperronienne et aurignacienne: étude de trois séries inédites de dents percées et comparaisons. C. R. Acad. Sci. (Ser. 2a) (Sci. Terre Planete/Earth Planet. Sci.) 325, 537–543 (1997).

46. D. Mylopotamitaki, et al., Comparing extraction method efficiency for high-throughput palaeoproteomic bone species identification. Sci. Rep. 13, 18345 (2023).

47. J. Cox, M. Mann, MaxQuant enables high peptide identification rates, individualized p.p.b.-range mass accuracies and proteome-wide protein quantification. Nat. Biotechnol. 26, 1367–1372 (2008).

48. Y. Perez-Riverol, et al., The PRIDE database resources in 2022: a hub for mass spectrometry-based proteomics evidences. Nucleic Acids Res. 50, D543–D552 (2022).

49. R Core Team, R: A Language and Environment for Statistical Computing. (2023). Available at: https://www.R-project.org/.

50. H. Wickham, et al., Welcome to the tidyverse. Journal of Open Source Software. (2019)

51. S. Firke, janitor: Simple Tools for Examining and Cleaning Dirty Data. (2023). Available at: https://CRAN.R-project.org/package=janitor.

52. A. Kassambara, ggpubr: “ggplot2” Based Publication Ready Plots. (2023). Available at: https://CRAN.R-project.org/package=ggpubr.

53. B. R. Mills, MetBrewer: Color Palettes Inspired by Works at the Metropolitan Museum of Art. (2022). Available at: https://CRAN.R-project.org/package=MetBrewer.

54. M. Mackie, et al., Palaeoproteomic Profiling of Conservation Layers on a 14th Century Italian Wall Painting. Angew. Chem. Int. Ed Engl. 57, 7369–7374 (2018).

55. A. Kuznetsova, P. B. Brockhoff, R. H. B. Christensen, lmerTest Package: Tests in Linear Mixed Effects Models. Journal of Statistical Software (2017).

56. D. Bates, M. Mächler, B. Bolker, S. Walker, Fitting Linear Mixed-Effects Models Using lme4. Journal of Statistical Software (2015).

57. W. N. Venables, B. D. Ripley, Modern Applied Statistics with S. (2002). Available at: http://www.stats.ox.ac.uk/pub/MASS4.

