## Supplementary Information for "New methods on the block: Taxonomic identification of archaeological bones in resin-embedded sediments through palaeoproteomics"

### Archaeological Context

For this study, we used archaeological resin-impregnated blocks from the Paleolithic sites of Bacho Kiro, La Ferrassie and Quinçay. The samples were taken as intact sediment blocks with the location of the resin-impregnated blocks presented in Table S1.

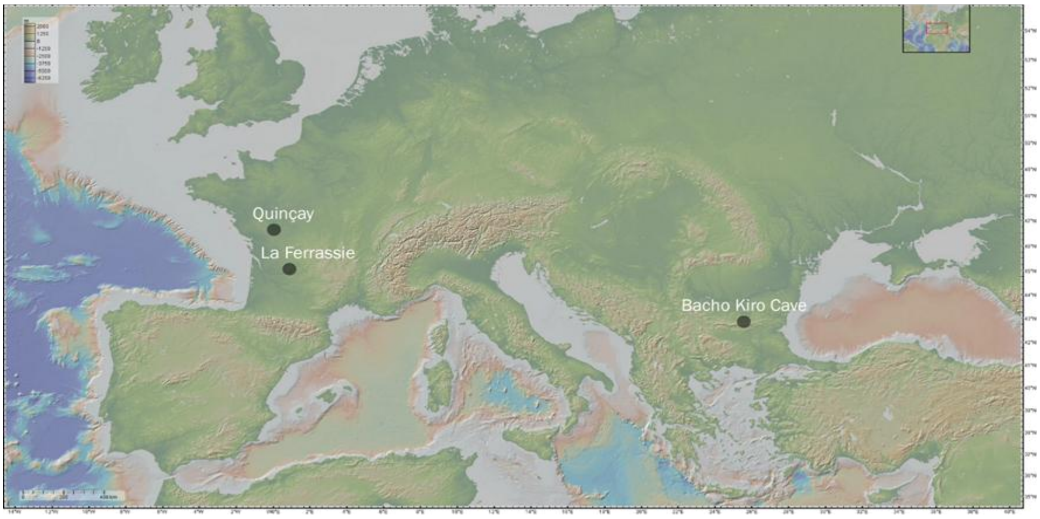


**Figure S1: Location of the studied archaeological sites within Europe.** Map produced from geomapapp.org

**La Ferrassie (Dordogne, France)**

Sample LAF 321: Location within site: Western sector; Layers 6-5b, coordinates: $X_{up}$=102.930; $Y_{up}$=111.419; $Z_{up}$=-1.068; $X_{low}$=102.984; $Y_{low}$=111.424; $Z_{low}$=-1.189

Impregnation protocol: Samples were extracted using either plaster of Paris bandages from exposed profiles at the site or soft paper. Samples were processed by Thomas Beckmann (Schwülper-Lagesbüttel, Germany) with dehydration and vacuum impregnation with polyester resin. The samples were trimmed to 9 cm x 6 cm sizes for the production of thin section slides.

Sample LAF 406.1: Location within site: Northern Sector; Layers E1, D and C. The block LAF406.1 only has layer E1; coordinates: $X_{up}$=117.165; $Y_{up}$=109.930; $Z_{up}$=-1.881, $X_{low}$=117.250; $Y_{low}$=109.907; $Z_{low}$=-2.253

Impregnation protocol: Samples were extracted using either plaster of Paris bandages from exposed profiles at the site or soft paper. Samples were processed by Thomas Beckmann (Schwülper-Lagesbüttel, Germany) with dehydration and vacuum impregnation with polyester resin. The samples were trimmed to 9 cm x 6 cm sizes for the production of thin section slides.

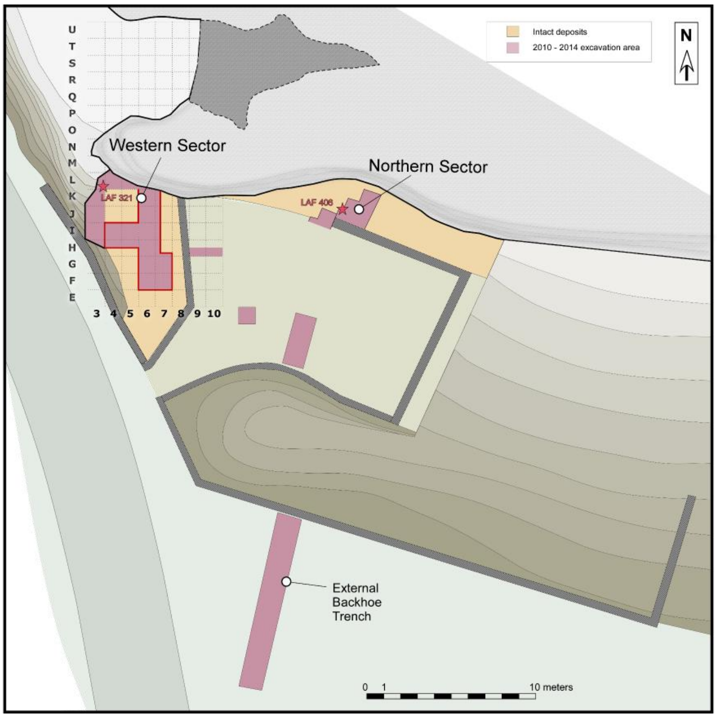


**Figure S2. La Ferrassie plan map.** Indicated are the approximate location of the resin-impregnated blocks LAF 321 in the Western Sector of excavations, and LAF 401 in the Northern Sector.

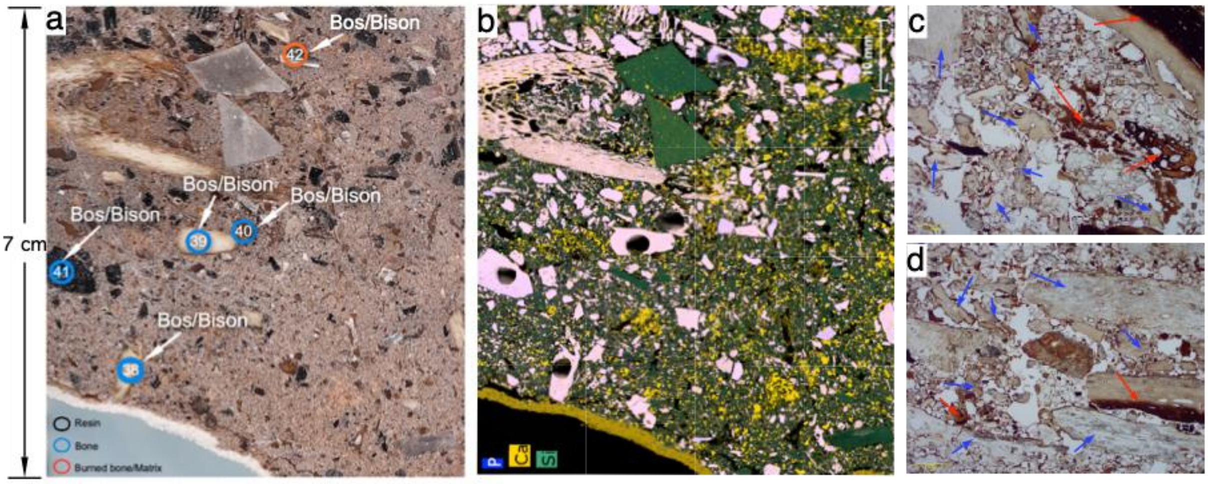


**Figure S3. Distribution of samples and their taxonomic attribution from La Ferrassie (resin-impregnated block LAF 321.2 side B)** Here, a) is the resin-impregnated block annotated with sampling locations and their taxonomic attribution; b) mXRF elemental mapping of phosphorus (P), calcium (Ca) and silicon (Si) showing the distribution of bones (pink), pure limestone (yellow) and silica associated with both sand-sized quartz and larger knapped flints (green); c) and d) are photomicrographs showing the ubiquitous presence of bones (blue arrows) and carbonized burned bones (red arrows) in these sediments, both images are taken in plane-polarized light and have a scale of 1 mm.

#### Bacho Kiro (Dryanovo, Bulgaria)

<u>Sample BK 501:</u> Location within site: Niche 1; Layer N1-I and N1-J, coordinates: Xup=92.679, Yup=96.884, Zup= -3.683, Xlow=92.63, Yellow=96.836, Zlow=-3.656

<u>Impregnation protocol:</u> Samples were extracted using either plaster of Paris bandages from exposed profiles at the site or soft paper. Samples were processed by Thomas Beckmann (Schwülper-Lagesbüttel, Germany) with dehydration and vacuum impregnation with polyester resin. The samples were trimmed to 9 cm x 6 cm sizes for the production of thin section slides.

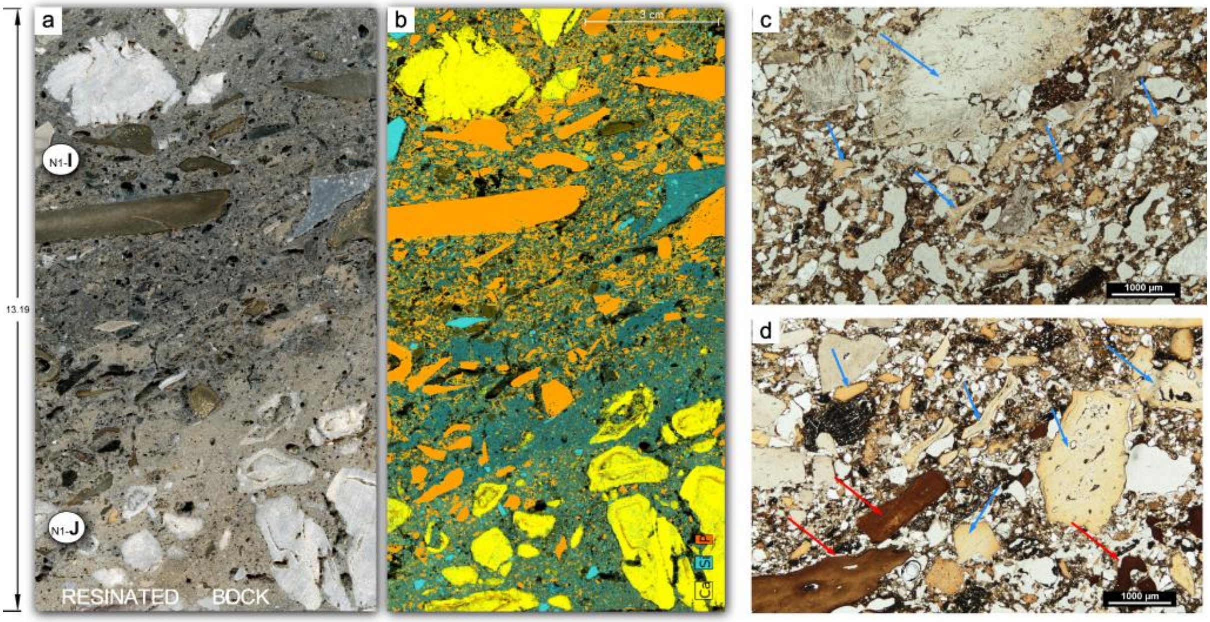


**Figure S4. Resin-impregnated block from Bacho Kiro (BK 501).** a) Photograph of the block with indication of archaeological layers represented in these deposits (N1-I and N1-J); b) mXRF elemental mapping showing the distribution of calcium in yellow, silicon in blue and phosphorous in red. Note the high concentration of bone fragments (with orange colorations) in these deposits, namely in the upper N1-I layer. Scale is 3 cm; c) and d) are photomicrographs in plane polarized light (PPL) showing the abundance of microscopic bone fragments in these deposits (blue arrows), including some carbonized bones (read arrows). Scale of both images is 1 mm.

**Quinçay (Poitiers, France)**

Sample GRL 103: Location within site: front1 profile; Layers 40 and 30b;

Impregnation protocol: Samples were extracted using either plaster of Paris bandages from exposed profiles at the site or soft paper. Samples were processed by Spectrum Petrographics (Vancouver, Washington, USA) with dehydration and vacuum impregnation with polyester resin. The samples were trimmed to 7.5 cm x 5 cm sizes for the production of thin section slides.

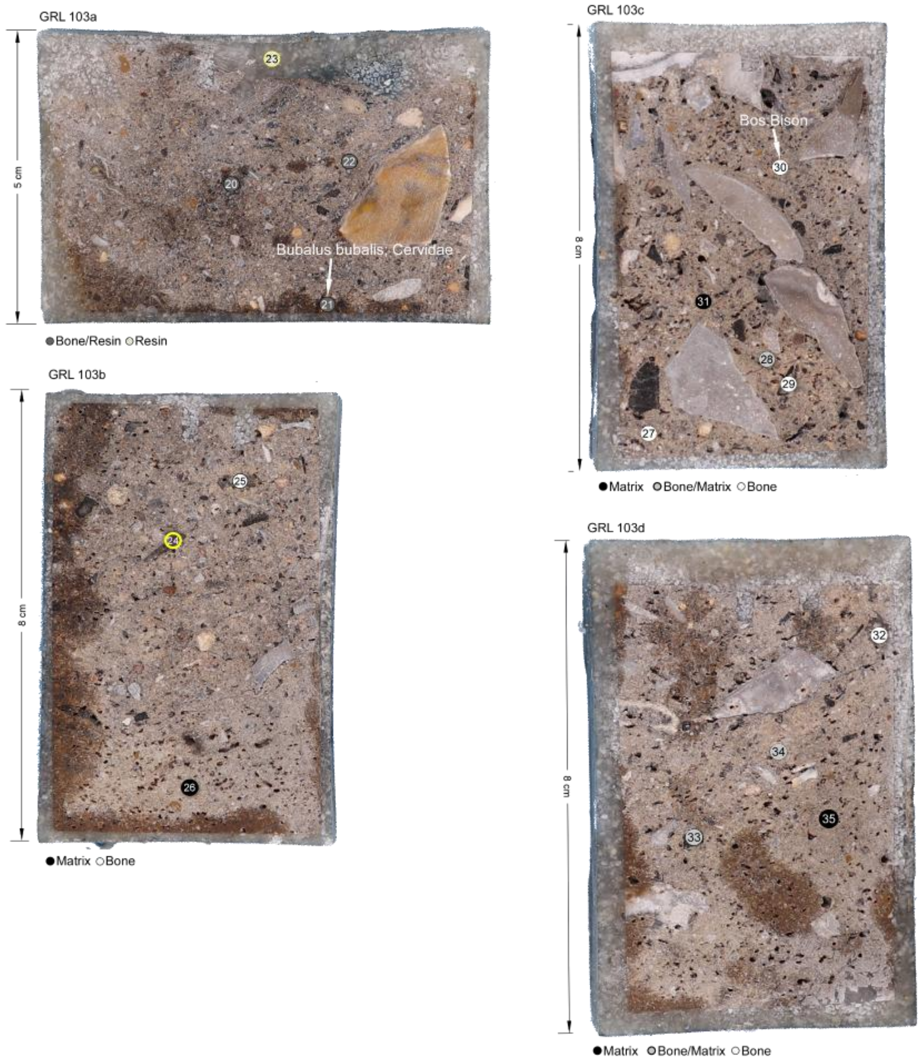


**Figure S5. Resin-impregnated block samples from the site of Quinçay.** The numbers represent the samples for proteomics with circles representing matrix (black), bone/matrix (gray) and bone (white).

**Table S1: Resin-impregnated blocks sampled for proteomic analysis.** Each resin-impregnated block and slab (face of the sampled block) have a unique identification. Sample number, amount of material sampled for protein extraction, and archaeological layer assignment are indicated.

| Site | Block | Slab & side | Sample # | Note | Weight (mg) | Archaeological layer(s) |
|---|---|---|---|---|---|---|
| **Bacho Kiro** | BK 501 | BK501.1 sideB | 1 | | 15 | N1-I |
| | | BK501.1 sideB | 2 | | 34 | N1-I |
| | | BK501.1 sideB | 3 | Bone and matrix | 14 | N1-I |
| | | BK501.1 sideB | 4 | | 20 | N1-I |
| | | BK501.1 sideB | 5 | Matrix | 28 | N1-I |
| | | BK501.1 sideB | 6 | Resin and matrix | 18 | -- |
| | | BK501.2 sideB | 7 | | 14 | N1-I |
| | | BK501.2 sideB | 8 | | 31 | N1-I |
| | | BK501.2 sideB | 9 | Bone and matrix | 29 | N1-I |
| | | BK501.2 sideB | 10 | Black bone (or charcoal?) | 9 | N1-I |
| | | BK501.2 sideB | 11 | Black bone | 12 | N1-I |
| | | BK501.2 sideB | 12 | Matrix | 27 | N1-I |
| | | BK501.2 sideB | 13 | Matrix | 29 | N1-J |
| | | BK501 sideB | 44 | Bone | 6 | N1-J |
| | | BK501 sideB | 45 | Bone - spongy bone | 48 | N1-J |
| | | BK501 sideB | 46 | Matrix - layer I | 48 | N1-I |
| | | BK501 sideB | 47 | Matrix - layer J | 36 | N1-J |
| **La Ferrassie** | LAF 406 | LAF406.1 | 14 | | 15 | E1-D |
| | | LAF406.1 | 15 | | 18 | E1-D |
| | | LAF406.1 | 16 | Possibly two bones sampled | 4 | E1-D |
| | | LAF406.1 | 17 | Black bone | 5 | E1-D |
| | | LAF406.1 | 18 | Matrix | 15 | E1 |
| | | LAF406.1 | 19 | Resin | 7 | -- |
| | LAF 321 | LAF321.2 side A | 36 | Bone - cortical section of bone | 7 | 5b |

| | | | | | | |
|---|---|---|---|---|---|---|
| | | LAF321.2 side A | 37 | Bone - spongy section of bone | 35 | 5b |
| | | LAF321.2 side B | 38 | Bone - yellow bone | 17 | 5b |
| | | LAF321.2 side B | 39 | Bone - yellow bone | 21 | 5b |
| | | LAF321.2 side B | 40 | Bone - brown bone | 29 | 5b |
| | | LAF321.2 side B | 41 | Bone - black bone | 9 | 5b |
| | | LAF321.2 side B | 42 | Bone - white bone - and matrix | 1 | 5b |
| | | LAF321.2 side B | 43 | Resin | 28 | -- |
| **Quinçay** | GRL 103 | GRL103a | 20 | Bone and resin | 16 | 30b |
| | | GRL103a | 21 | Bone and resin | 17 | 30b |
| | | GRL103a | 22 | Bone and resin | 6 | 30b |
| | | GRL103a | 23 | Resin | 17 | -- |
| | | GRL103b | 24 | Bone | 1 | 30b |
| | | GRL103b | 25 | Bone? | 25 | 30b |
| | | GRL103b | 26 | Matrix | 9 | 40 |
| | | GRL103c | 27 | Bone | 12 | 30b |
| | | GRL103c | 28 | Bones | 7 | 30b |
| | | GRL103c | 29 | Bone | 29 | 30b |
| | | GRL103c | 30 | Bone and matrix | 6 | 30b |
| | | GRL103c | 31 | Matrix | 14 | 30b |
| | | GRL103d | 32 | Black bone | 8 | 30b |
| | | GRL103d | 33 | Bone and matrix | 18 | 30b-40 |
| | | GRL103d | 34 | Bone and matrix | 32 | 30b |
| | | GRL103d | 35 | Matrix | 24 | 30b-40 |

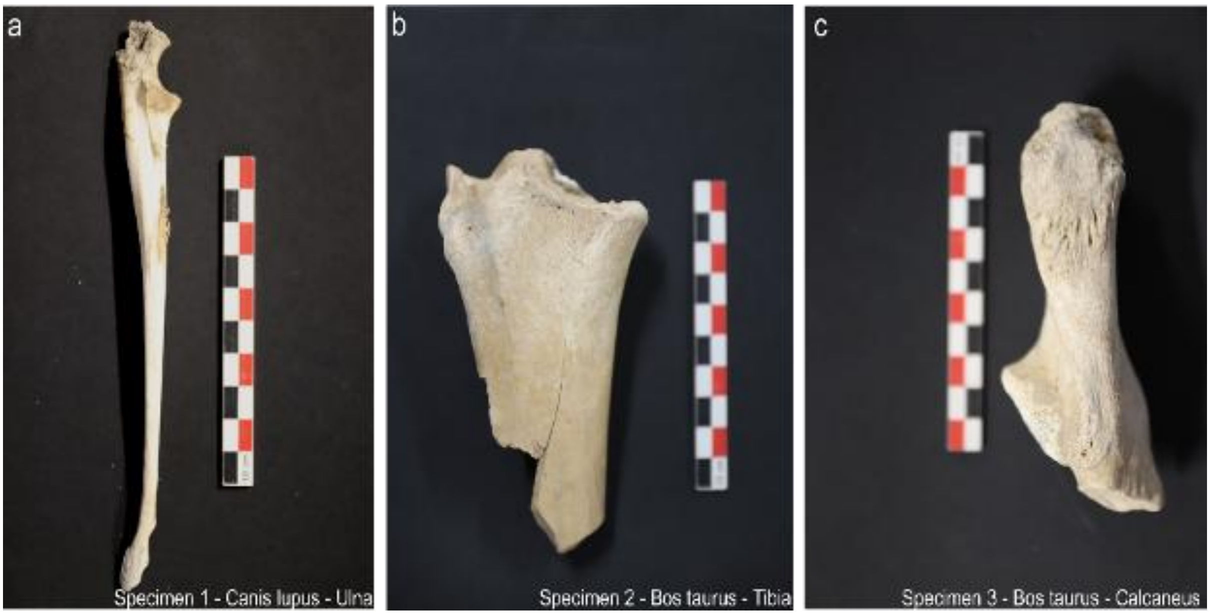


**Figure S6: Modern bone samples used prior to sampling.** Scales are 10 cm.

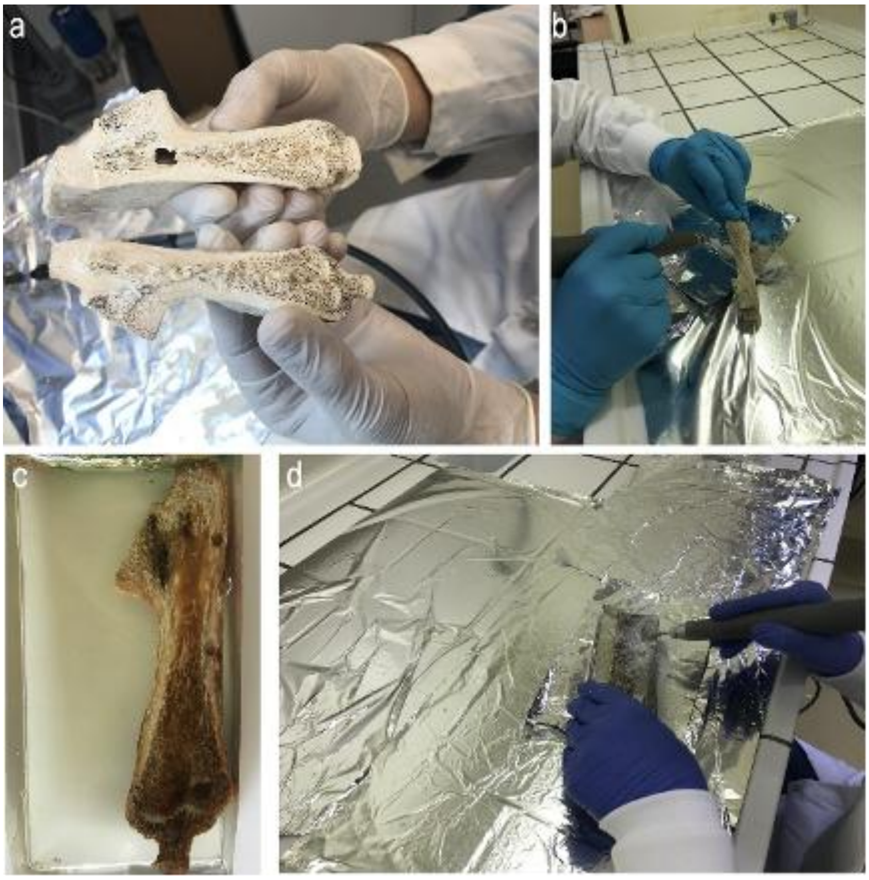


**Figure S7. Workflow for sampling modern bones.** Cutting of bones to expose internal structure (a), sampling (b), impregnation with resin (c) and sampling after resin impregnation (d).

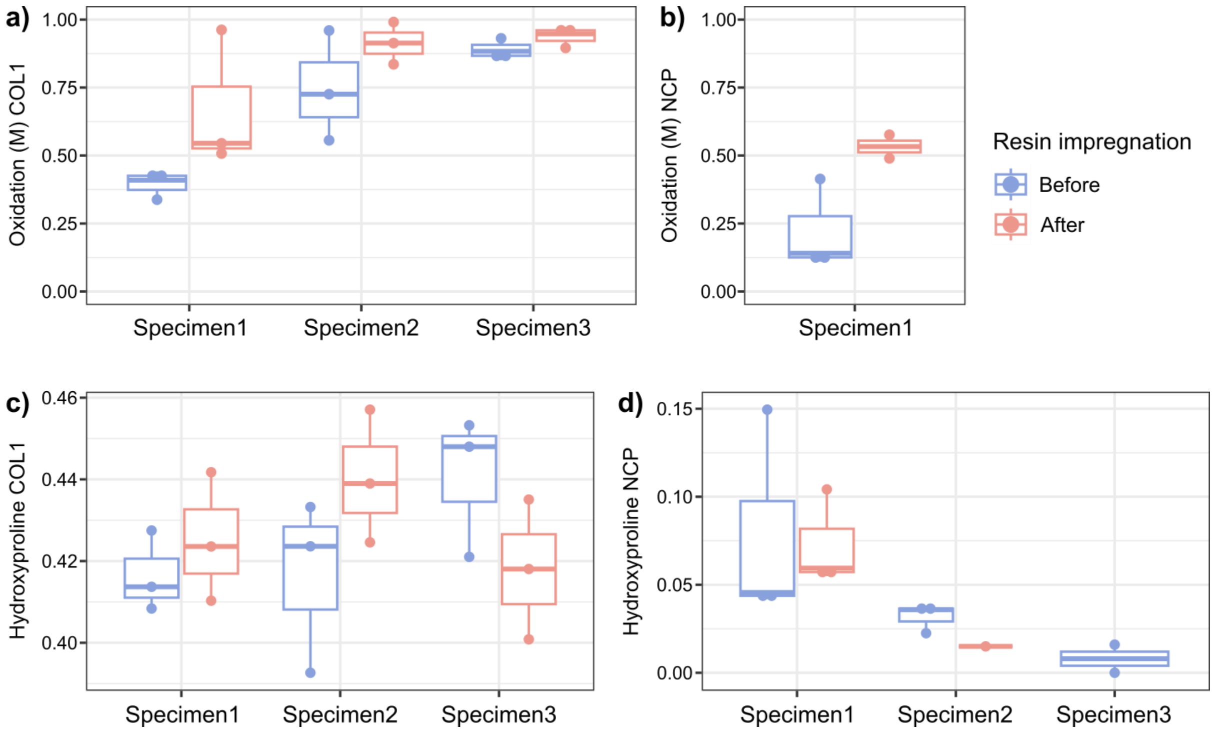


**Figure S8. Oxidation-related post-translational modifications for different protein groups.** a) Oxidation of methionine for COL1, b) Oxidation of methionine for NCPs, c) Hydroxyproline for COL1, d) Hydroxyproline for NCPs. Note the differences in the y-axis scale between the four panels. Only specimens containing more than 10 peptides with the amino acid of interest are included for each analysis.

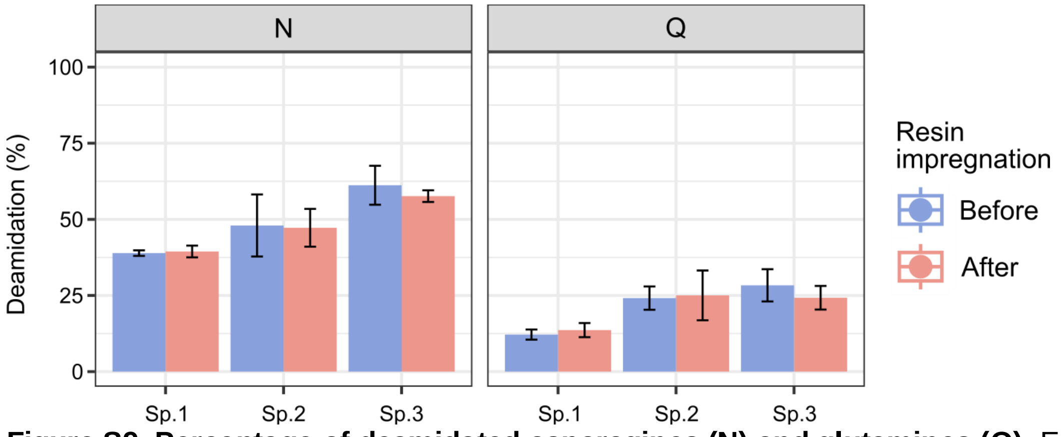


**Figure S9. Percentage of deamidated asparagines (N) and glutamines (Q).** Error bars indicate ± 1 standard deviation of mean deamidation of the three replicates.

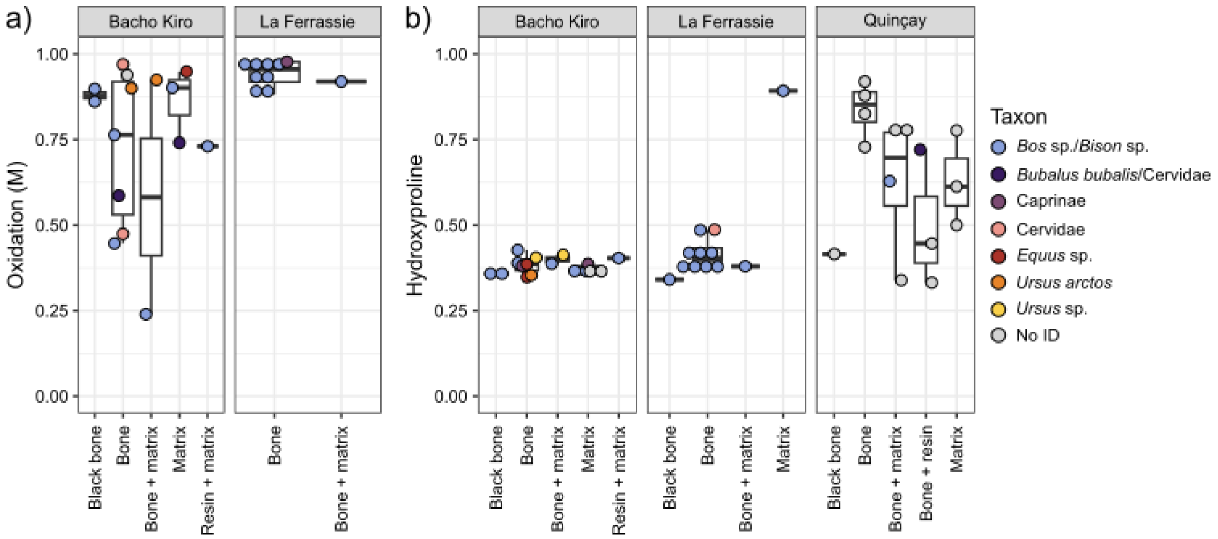


**Figure S10. Post-translational modifications of archaeological block samples.** a) Fraction of oxidated M, and b) Fraction of hydroxylated prolines. Each point is a sample, colored by taxonomic identification through SPIN. Only samples with >10 peptides containing the amino acid of interest are included.

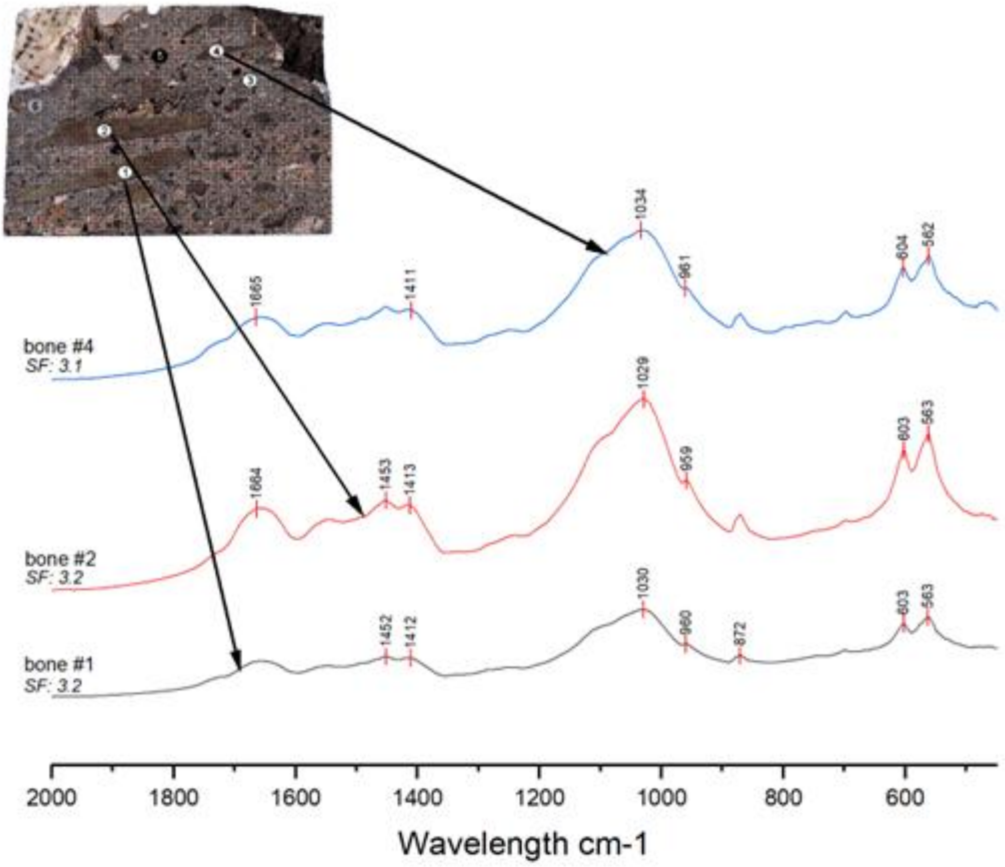


**Figure S11: Infrared spectra from bones in sample BK 501-1.** All specimens have absorption peaks compatible with unburned bones, with the presence of major phosphate attributed to PO4 absorption peak around 1,033 cm-1, the presence of the carbonate peaks at 872 cm-1 and 1415 cm-1, and the absence of the phosphate high temperature peak associated with OH libration band at ca. 630 cm-1 (1–4). SF: splitting factor.

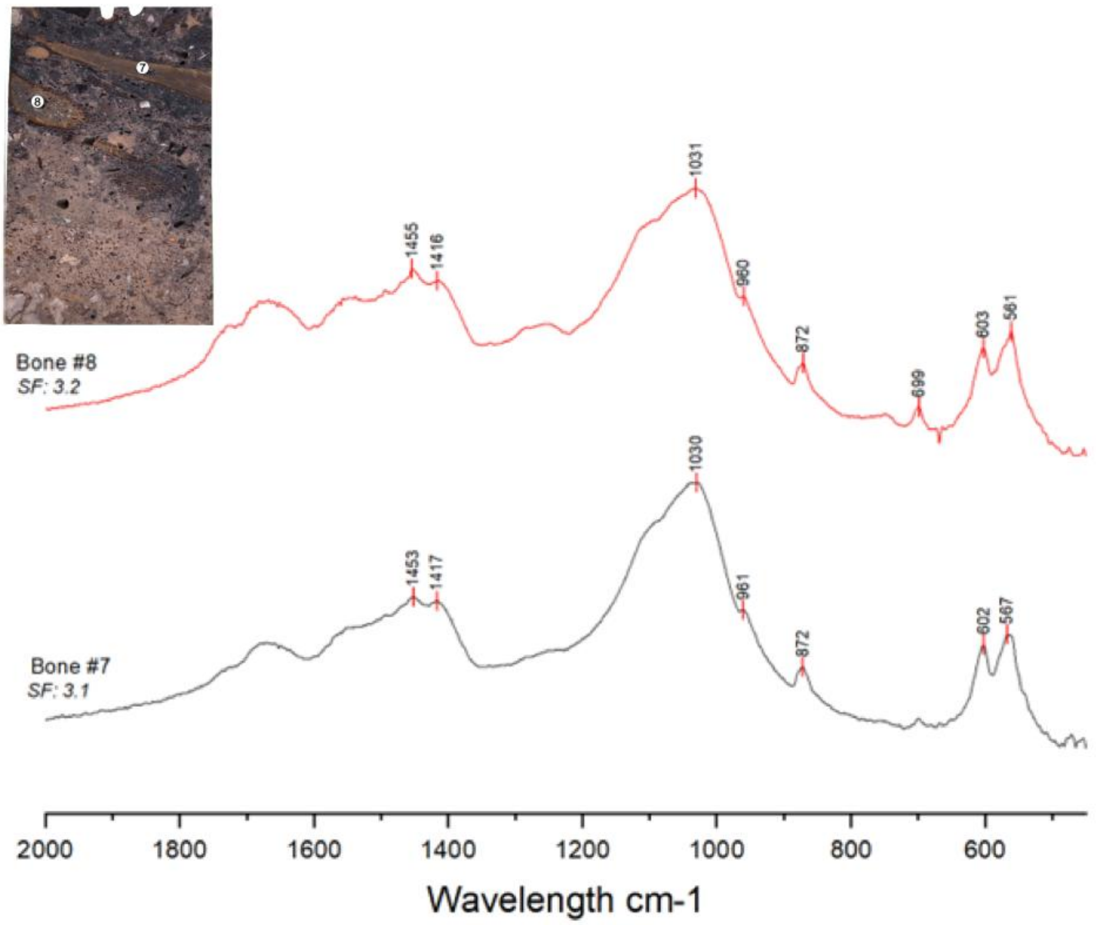


**Figure S12: Infrared spectra from bones in sample BK 501-2, showing non-calcined bones.**

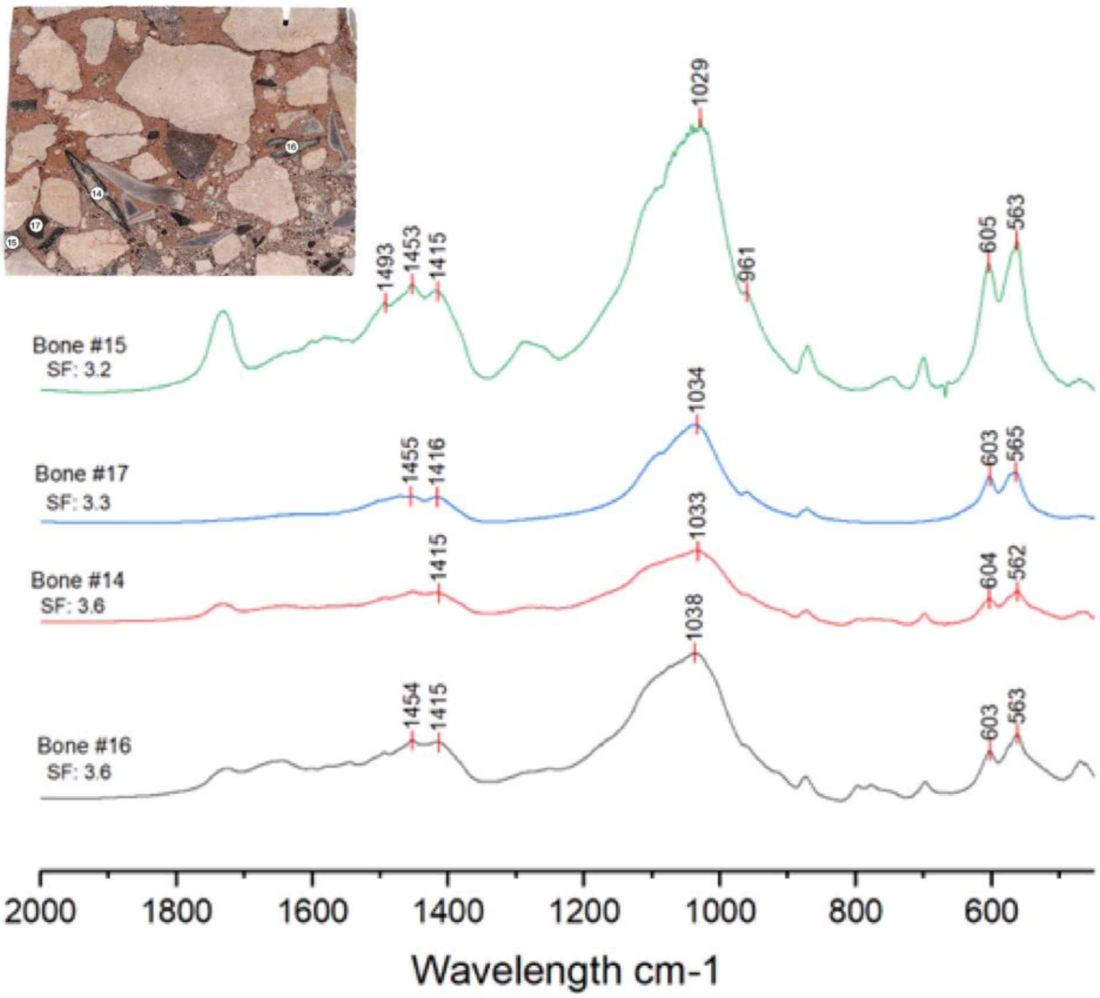


**Figure S13: Infrared spectra from bones in sample LAF 406-1, showing non-calcined bones.**

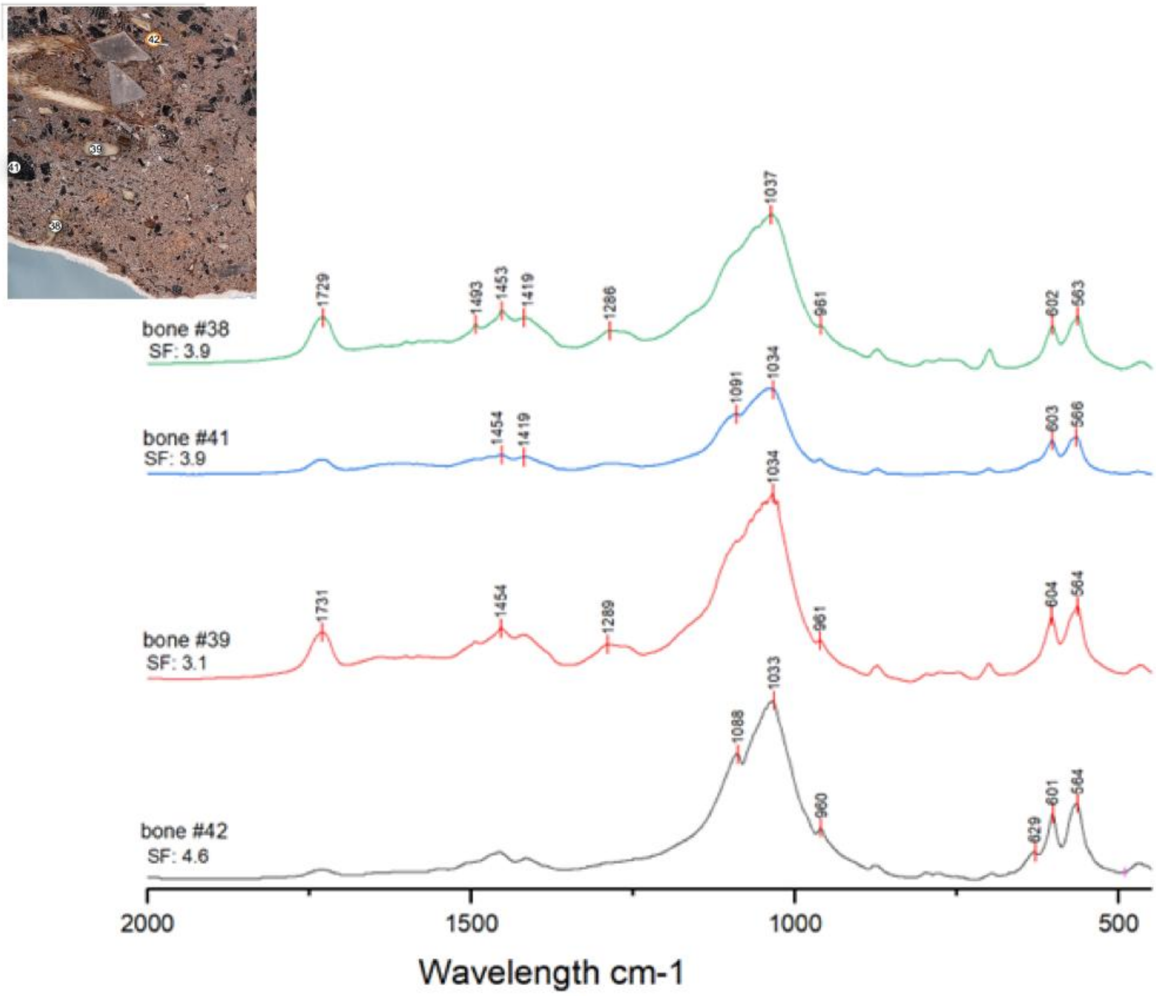


**Figure S14: Infrared spectra from bones in sample BK321-2.** Note the presence of the phosphate high temperature peak at ~630 cm-1 and the high splitting factor of 4.6 for sample bone #42 indicating calcination at a temperature above 700ºC. Note also the slight shoulder at 630 cm-1 present in same bone #41 associated with a black bone, indicating possible carbonization but at temperatures below 700ºC.

### References

1. C. L. Shaw, An evaluation of the infrared 630 cm−1 OH libration band in bone mineral as evidence of fire in the archaeological record. *J. Archaeol. Sci. Rep.* **46**, 103655 (2022).

2. M. C. Stiner, S. L. Kuhn, S. Weiner, O. Bar-Yosef, Differential burning, recrystallization, and fragmentation of archaeological bone. *J. Archaeol. Sci.* **22**, 223–237 (1995).

3. T. J. U. Thompson, M. Gauthier, M. Islam, The application of a new method of Fourier Transform Infrared Spectroscopy to the analysis of burned bone. *J. Archaeol. Sci.* **36**, 910–914 (2009).

4. S. Weiner, *Microarchaeology: Beyond the visible archaeological record* (Cambridge University Press, 2010).
